# Conformational dynamics associated with remote residues regulate the kinetic properties of homologous glutamate dehydrogenases (GDHs)

**DOI:** 10.1101/2024.04.12.589225

**Authors:** Barsa Kanchan Jyotshna Godsora, Parijat Das, Anjali Sairaman, Prasoon Kumar Mishra, Sandip Kaledhonkar, Narayan S. Punekar, Prasenjit Bhaumik

## Abstract

Glutamate dehydrogenase (GDH) is a key enzyme in all living organisms and some of the GDHs exhibit substrate-dependent homotropic cooperativity. However, the mode of allosteric communication during the homotropic effect in GDHs remains poorly understood. In this study, we examined two homologous GDHs, *Aspergillus niger* GDH (AnGDH) and *Aspergillus terreus* GDH (AtGDH), with differing substrate utilization kinetics to uncover the factors driving their distinct behavior. The crystal structures and first-ever cryo-EM structures of apo-AtGDH captured arrays of conformational ensembles. Comparative structural analysis has revealed a wider mouth opening in allosteric AnGDH (∼ 21 Å) compared to non-allosteric AtGDH (∼17 Å) in their apo states. A network of interaction related to the amino acid substitutions in Domain II is responsible for differential structural dynamics in these GDHs. Remarkably, we identified one remotely located substitution in Domain II, i.e., R246 to S, a part of the network, which reversed the kinetic properties of AtGDH into an allosteric one and controls the mouth opening. Our data also indicate that dynamic discrepancy influences the substrate binding affinity and catalytic activity in AnGDH and AtGDH. We have successfully demonstrated for the first time, that remotely located residues and the conformational dynamics regulate the kinetic properties in homologous GDHs.

## Introduction

Enzymes are biological catalysts that are involved in numerous biological processes. The molecular basis of the mechanism of how enzymes enhance the rate of a reaction and regulate their catalytic efficiency has remained unclear despite several studies in this field for decades. It is even more puzzling that the homologous enzymes, which share conserved structural fold and active site residues, exhibit disparities in their kinetic properties, such as substrate affinity, turnover, temperature adaptation, allosteric regulation, etc. (Bae & Philips, 2004; Kamata et al., 2004; Torgeson et al., 2022).

Studies on homologous monomeric proteins highlighted the role of remotely located amino acid substitutions far from the active site for their varied kinetic properties (Bhabha et al., 2011). Substitutions on the surface residues alter the intrinsic dynamics (e.g., dihydrofolate reductase DHFR) and conformational dynamics (e.g., glucokinase), thereby affecting the kinetic properties of an enzyme. Moreover, conformation differences in the apo form of homotetrameric D-lactate dehydrogenase (D-LDH) from *Fusobacterium nucleatum* (FnLDH) and *Pseudomonas aeruginosa* (PsLDH) have affected the inter-subunit packing, leading to positive cooperativity in FnLDH while negative cooperativity in PsLDH for the same substrate pyruvate (Furukawa et al., 2018). Several homologous monomeric proteins like adenylate kinase (Bae & Philips, 2004), glucokinase (Kamata et al., 2004), etc., have been well studied, while the knowledge of oligomeric proteins from the same perspective is scanty. A hexameric glutamate dehydrogenase (GDH) across the different kingdoms exhibits various modes of allosteric regulation (Wang & Engel, 1995; Noor & Punekar, 2005; DeLuna et al., 2001; Smith et al., 2002). Here, we have considered studying GDH to decipher the molecular mechanism that controls kinetic properties and allosteric regulation in this enzyme.

GDH is an important metabolic enzyme that catalyzes the reversible conversion of α-ketoglutarate (AKG) to L-glutamate. Being at a branch point between the carbon and nitrogen cycles, it is involved in several downstream biological processes (Lee et al., 2012; Mathioudakis et al., 2019; Lee et al., 2019) and consequently, linked to several metabolic (e.g., tumorogenesis and PAcidemia) (Roginski et al., 2019) and neural diseases (Parkinson’s and Alzheimer’s) (Smith et al., 2019; Basith et al., 2021). Systematic investigations of the GDHs promise to broaden our understanding of this widely recognized phenomenon.

We have considered a promiscuous homologous fungal enzyme system to study the substrate-dependent homotropic cooperativity in lower eukaryotic GDHs. Among *Aspergilli* spp., *Aspergillus niger* GDH (AnGDH) and *Aspergillus terreus* GDH (AtGDH) share 88 % amino acid sequence identity, yet disparities exist in several kinetic properties such as substrate affinity, isophthalate inhibition, mode of substrate AKG utilization, etc. (Choudhury & Punekar, 2007; Agarwal et al., 2019). AnGDH exhibits homotropic cooperativity toward AKG substrate saturation (*S*0.5 = 5.6 mM, *n*H = 2.9), while AtGDH shows a Michaelis-Menten kinetics for the same (*K*m = 4.3 mM). In our study, we have focussed on deciphering the molecular mechanism that controls one of the kinetic differences, i.e., substrate-dependent homotropic cooperativity in these homologous *Aspergilli* GDHs.

Despite structural and biochemical studies (Prakash et al., 2018) on AnGDH, the atomistic details of the network that regulates the homotropic cooperativity for AKG substrate binding are unknown. Furthermore, substantial experimental evidence that supports the intrinsic dynamic nature of GDHs is missing. Therefore, determining more GDH structures with different conformational states and their analysis and complementary biochemical characterizations would help to identify the apo-state conformers that influence the kinetics of substrate binding. Moreover, comparative structural analysis of these homologous enzymes -AnGDH and AtGDH, will help to unravel the underlying mechanistic details of homotropic cooperativity.

Our study presents the first cryo-EM structures of fungal GDHs. Here, we report the structures of multiple conformational ensembles of apo AtGDH determined using X-ray crystallography and cryo-Electron Microscopy (cryo-EM) revealing the inherent dynamicity in GDHs. Based on structural and kinetic studies on AtGDH and AnGDH, we have identified the key substitution responsible for inducing kinetic disparities in these homologous enzymes. Henceforth, we propose for the first time that the intrinsic conformational dynamics of GDHs is responsible for their kinetic regulation. Thus, this study highlights the significance of surface exposed remote residues in controlling the protein dynamics and kinetic behavior in an enzyme.

## Material and Methods

### Expression constructs of wild type (WT) AnGDH and AtGDH

The expression construct (without His_6_ tag) of *Aspergillus niger* NADP^+^-dependent GDH was available in our lab (Prakash et al., 2018). Using the above plasmid construct, the *gdh*A gene was amplified using Q5 Polymerase (NEB) and the primers (Table S5). The amplified product comprised flanking restriction sites at either end, TEV protease cleavage site (ENLYFQG), and *gdh*A cds between them. The gene construct was inserted into pET28a (+) vector between the restriction site *Nde*I and *Xho*III (ThermoFischer) to include N-terminal His_6_ tag. The plasmid construct was transformed in *E. coli* DH5α cells. The expression plasmid construct confirmed by sequencing was then transformed into Δ*gdh E. coli* BL21 (DE3) cells. For AtGDH, the construct mentioned previously (Godsora et al., 2022) was used.

### Site-directed mutagenesis

The plasmid containing the *Atgdh* gene was used as a template for mutating amino acids at the desired positions. The high-fidelity Q5 Polymerase enzyme (NEB) was used to amplify the whole plasmid length using the specific site-directed mutagenesis (SDM) primers listed below (Table S5). AtGDH SDM mutants have been generated sequentially using the previous mutation background as the template base.

Following the NEB deletion mutagenesis protocol, the first deletion mutation in the *Atgdh* gene were made at positions - T262 and A263. The *Atgdh*-containing plasmid was amplified using the primers Del_Atgdh-FP and Del_Atgdh-RP (Table S5), which lacked the residues T262-A263. It was followed by treatment of the amplicon with T4 Polynucleotide kinase (Thermo Scientific) for 1 h at 37 ℃. Further, the linear amplicon was ligated using the T4 Ligase enzyme for 4 h at 37 ℃ (Thermo Scientific) and then followed by heat-inactivation at 65 ℃ for 10 min. Lastly, it was exposed to digestion with *Dpn*I Fast digest enzyme (Thermo Fischer) for 3 h at 37 ℃ to remove the parental strands. The treated amplicon was finally transformed into the *E. coli* DH5α cells for maintaining the plasmids. The mutation was confirmed by gene sequencing (First Base sequencing, Malaysia). The confirmed plasmid construct was then transformed in Δ*gdh E. coli* BL21 (DE3) expression host cells. The confirmed plasmid construct and the expressed protein are referred to as 2^nd^_AtGDH-ΔT262-A263.

The second mutation consisted of four amino acid substitutions at positions-K260, D261, K264, and D265 simultaneously. It is located next to the insertion site (T262-A263) in the *Atgdh* gene. The 2^nd^_AtGDH-ΔT262-A263 plasmid was used as the template to mutate the residues-K260N, D261G, K264E, and D265G, using the ATGDH-STEP2-FP forward and ATGDH-STEP2-RP reverse primers listed in Table S5 As described above, the amplicon was treated stepwise with T4 Polynucleotide kinase, T4 Ligase, and *Dpn*I. The mutation was confirmed by gene sequencing. The confirmed construct containing plasmid and the expressed protein is referred to as 6^th^_AtGDH-KDTAKD. It was followed by transforming the confirmed plasmid construct 6^th^_AtGDH-KDTAKD into Δ*gdh E. coli* BL21 (DE3) cells.

Further, the other mutants (R246S and A216Q) were generated, keeping the previous background as the template for mutating the amino acids at the desired positions in the *Atgdh* gene. The plasmid constructs and the expressed proteins are designated as 7^th^_AtGDH-R246S and 8^th^_AtGDH-A216Q.

In the case of AnGDH, single mutations at positions - Q216A and S246R were made. The plasmid containing the *Angdh* gene was used as the template. The high-fidelity Q5 Polymerase enzyme (NEB) was used for each mutagenesis to amplify the whole plasmid length using the specific SDM primers listed below (Table S5). The plasmid constructs and the expressed proteins are designated as AnGDH-Q216A and AnGDH-S246R.

All the mentioned amplicons were digested by restriction enzyme *Dpn*I (Thermo Fischer) at 37 ℃ for 4 h to remove the parental plasmid. Further, the treated amplicons were individually transformed into the *E. coli* DH5α cells for storage. The mutations were further confirmed by gene sequencing. Then, the confirmed constructs were transformed in Δ*gdh E. coli* BL21 (DE3) cells for protein expression.

### Expression and purification of WT AtGDH and AnGDH

Recombinant WT AtGDH was expressed, and Ni-NTA affinity and size exclusion chromatography were used to purify the enzyme as described previously (Godsora et al., 2022). The cell containing the plasmid bearing *Angdh* construct was inoculated in 5 ml LB broth containing kanamycin (50 µg/ml) overnight. The overnight grown cultures were re-inoculated in fresh 500 ml LB broth containing kanamycin (50 µg/ml) and grown at 37 ℃ till it reached an optical density (OD) of 0.7–0.8 at 600 nm. Further, the cells were induced with 0.4 mM IPTG and grown in slow shaking for another 12 h at 22 ℃. The same protocol was used for cell harvesting and disruption as described for WT AtGDH (Godsora et al., 2022). Further, two rounds of Ni-NTA purifications (without and with TEV digestion performed) were conducted, as mentioned for WT AtGDH (Godsora et al., 2022), to remove the His_6_ tag. Final purification was performed with size exclusion chromatography using a Superdex 200 pg 16/600 gel filtration column (120 ml bed volume, GE Healthcare). At every purification step, the purity of AtGDH or AnGDH was analyzed using the 12% SDS-PAGE, and protein quantification was done using the standard Bradford assay.

### Expression and purification of the AtGDH and AnGDH mutants

All the confirmed clones of AtGDH mutants (designated as 2^nd^_AtGDH-ΔT262-A263, 6^th^_AtGDH-KDTAKD, 7^th^_AtGDH-R246S, and 8^th^_AtGDH-A216Q) and AnGDH mutants (AnGDH-Q216A and AnGDH-S246R) were expressed and purified in a similar way as described for wild types - AtGDH (Godsora et al., 2022) and AnGDH.

Both the AtGDH and AnGDH mutants were purified using the Ni-NTA batch method. Initially, the 2 ml Ni-NTA resin (cOmplete His-tag purification resin, Roche) was washed with 10 column volume (CV) of 20% ethanol, followed by 10 CV wash with MiliQ and then equilibrated with 20 CV of equilibration buffer (EB) (30 mM Phosphate buffer pH 7.5). The filtered supernatant was loaded on the equilibrated Ni-NTA resin and incubated for 1 h with continuous slow shaking at 4 °C to facilitate protein binding. The Ni-NTA resin with bound protein was washed separately with 25 mM and 50 mM. The elution was performed using 125 mM, 250 mM, and 500 mM imidazole containing EB buffer pH 7.5. The purity of the eluted protein sample was analyzed using 12% SDS-PAGE.

The eluted AtGDH and AnGDH mutants were treated separately with the TEV protease to remove the His_6_ tag, as mentioned (Godsora et al., 2022). Further, to separate the cleaved and non-cleaved His_6_-tagged proteins, the second round of Ni-NTA purification (batch method) was performed. Next, size exclusion chromatography (Superdex 200 pg 16/600 column, 120 ml bed volume, GE Healthcare) was performed as a final purification step. After every purification step, the purity of the desired AtGDH and AnGDH mutant proteins was checked using the 12% SDS-PAGE. The protein estimation was done using the standard Bradford assay.

### Circular Dichroism (CD) of AtGDH and AnGDH

CD spectral scans at near UV (198-260 nm) were performed at room temperature for the wild types-AnGDH and AtGDH, and their mutants in the CD spectropolarimeter (Model-J, Jasco, Japan). The spectral bandwidth was kept at 5 nm. Baseline correction was done using the respective buffer. The sample concentration of 5 μM was considered, and the experiment was done in triplicates. Data smoothening was done in Jasco Spectra Manager, and then data were plotted in SigmaPlot.

### Enzyme activity assays and kinetic characterizations

The NADP^+^-dependent GDH catalyzes the reductive amination of α-ketoglutarate to L-glutamate in the presence of excess ammonia and coenzyme NADPH, termed forward activity. The forward activity of AtGDH and AnGDH was measured using the method described earlier (Noor & Punekar, 2005; Prakash et al., 2018; Agarwal et al., 2019; Godsora et al., 2022).

In the substrate saturation kinetics, the standard activity assay was modified, where at once the concentration of one substrate was varied while fixing the other substrates at their saturating concentration (10 mM α-ketoglutarate, 10 mM NH_4_Cl, 0.1 mM NADPH). A varying concentration of α-ketoglutarate in the 0–50 mM (adjusted to pH 8.0) was used. The enzyme assay was performed at room temperature in 1 ml reaction volume where 600 µl cocktail pH 8.0 (cocktail contained 100 mM Tris-Cl and 10 mM NH_4_Cl, adjusted with MQ to make 60 ml total volume), a fixed volume of 0.1 mM NADPH (∼0.622 *A*_340nm_), purified enzyme and varying concentration of α-ketoglutarate were used. The enzyme volume was adjusted to attain below 10% substrate conversion to capture the initial velocities. The experimental values were fitted to suitable rate equations such as Michaelis-Menten, Uncompetitive, or Hill. The non-linear regression curve in SigmaPlot 12.0 software plotted the data. The data points represent the calculated rate velocities, whereas the line represents the best fit to the curve. The experiments were performed in triplicates at room temperature.

### Crystallization of AtGDH and its mutants

A sitting or hanging drop vapor diffusion method was used to set up the crystallization of the apo-AtGDH, AtGDH-AKG bound binary complex, 2^nd^_AtGDH-ΔT262-A263 mutant, and 7^th^_AtGDH-R246S mutant-AKG bound binary complex. Several commercially available crystallization screens such as JCSG-plus (Molecular Dimensions), PEG suite (Qiagen), PEG Rx (Hampton Research), PEG/Ion (Hampton Research), and Index (Hampton Research) were used to obtain the initial crystal hits. The crystallization drops (0.6 μl) containing 0.3 μl of pure WT AtGDH (in apo or binary complex form) or its mutants and 0.3 μl of mother liquor were mixed in 1:1 or 1:2 ratios and equilibrated against the 50 μl reservoir containing mother liquor. The screens were set up in the crystallization trays using the automated Phoenix robot (Art Robins) at the Protein Crystallography Facility, Indian Institute of Technology Bombay. The crystallization trays were incubated at 22 ℃ in a vibration-free incubator. The crystal growth was frequently monitored under the stereomicroscope.

To obtain the crystals of the AtGDH binary complex with substrate AKG, the pure AtGDH (16 mg/ml or 320 μM) and substrate AKG (final concentration 6 mM) were pre-incubated for 1 h at 4 ℃. Several hits for the crystals of AtGDH-AKG binary complex were obtained in the PEG Rx, PEG suite, Index, and PEG Ion screens. Well diffracting crystals of the AtGDH-AKG binary complex appeared in 0.2 M ammonium acetate, 0.1 M Tris pH 8.5, 25% w/v polyethylene glycol 3,350 appeared after a year. Later, on solving the structure of the AtGDH-AKG binary complex, no electron density for the substrate AKG was observed, so hereafter, it is considered as apo form and referred to as AtGDH-I.

Further, the initial crystal hit for the apo form AtGDH (20 mg/ml) was obtained in 0.2 M MgCl_2_, 0.1 M Tris pH 7, and 10% PEG 8000. The hanging drop method was used for improving the quality of the crystals by varying the drop sizes and protein concentrations. AtGDH (0.5 μl) and mother liquor (1.5 μl) were mixed in a 1:4 ratio and equilibrated against the 0.5 ml mother liquor serving as a reservoir. The best diffracting quality apo-AtGDH crystal appeared in a week and grew to maximum size within three weeks. Here on, the crystal of apo form AtGDH is designated as AtGDH-II.

The hanging drop vapor diffusion method was used to crystallize the apo form 2^nd^_AtGDH-ΔT262-A263 and 7^th^_AtGDH-R246S-AKG bound binary complex. The good diffracting crystal of 2^nd^_AtGDH-ΔT262-A263 mutant (13 mg/ml) appeared in the mother liquor containing 15% PEG3350, 0.1 M HEPES pH 7.0, 0.2 M NaCl condition in a drop size (1:1) after two weeks.

To obtain the crystals of the 7^th^_AtGDH-R246S binary complex with substrate AKG, the pure 7^th^_AtGDH-R246S (15 mg/ml) and substrate AKG (final concentration 100 mM) were pre-incubated overnight at 4 ℃. A well diffracting crystal for the 7^th^_AtGDH-R246S binary complex appeared in 0.2 M Sodium thiocyanate, 20% PEG 3350, in drop size of 1:2 ratio in a week and grew to maximum size in two weeks. On solving the 7^th^_AtGDH-R246S-AKG binary complex structure, no electron density for the substrate AKG was observed, so therefore, it is considered as apo form.

### X-ray diffraction, data collection, and processing

Several cryoprotectants, such as glycerol, MPD, and ethylene glycol (10 – 40%), were screened against the nitrogen stream (∼100 K). The best cryo-protectant for AtGDH-I was 25% glycerol, 30% PEG 400 for 2^nd^_AtGDH-ΔT262-A263, while 30% and 40% MPD served as cryoprotectant for AtGDH-II and 7^th^_AtGDH-R246S crystals, respectively. All the diffraction data sets were collected from the frozen crystals using the rotation method. For each case, a single crystal was harvested using a nylon cryo-loop from the respective drop. It was immersed in their respective cryoprotectants and flash-frozen to a liquid nitrogen stream at 100 K. The diffraction spots of AtGDH-I, AtGDH-II, and 2^nd^_AtGDH-ΔT262-A263 were collected by rotating method at the home source using the CuKα X-ray radiation, which is generated by the Rigaku Micromax 007HF X-ray generator fitted with a Rigaku R-Axis IV++ detector (Protein Crystallography Facility, IIT Bombay). The crystal of 7^th^_AtGDH-R246S was briefly transferred to cryoprotectant and then immediately flash-frozen in liquid nitrogen for storage. The frozen crystals of 7^th^_AtGDH-R246S were then transferred to a liquid nitrogen stream at 100 K for data collection. The diffraction was performed at the PX-BL21 beamline of the Indus 2 synchrotron, RRCAT, Indore, India. The diffraction images of 7^th^_AtGDH-R246S were collected at 1° oscillations at 0.9794 Å wavelength on the attached Rayonix MX225 CCD detector.

The diffraction data sets of AtGDH-I, AtGDH-II, 2^nd^_AtGDH-ΔT262-A263, and 7^th^_AtGDH-R246S were processed (indexing, integration, and scaling) using XDS software (Kabsch, 2010). Unless mentioned, the default parameters were considered for processing the diffraction data sets. Finally, the recorded intensities from the data sets of AtGDH-I, AtGDH-II, 2^nd^_AtGDH-ΔT262-A263, and 7^th^_AtGDH-R246S were converted to structure factor by the F2MTZ and CAD program of CCP4 (Winn et al., 2011). The data collection statistics are presented in Table S1.

### Crystal structure determination and refinement

The initial phases were obtained by the molecular replacement (MR) method. The calculation of the Matthews coefficient (*V*_M_) (Matthews, 1968) for the AtGDH-I, AtGDH-II, 2^nd^_AtGDH-ΔT262-A263, and 7^th^_AtGDH-R246S crystals were 3.86 Å^3^ Da^-1^ (68% solvent content), 2.72 Å^3^ Da^-1^ (54.78% solvent content), 3.86 Å^3^ Da^-1^ (68% solvent content) and 2.50 Å^3^ Da^-1^ (50.8% solvent content), respectively. The asymmetric units were predicted to have two, six, two, and two molecules for AtGDH-I, AtGDH-II, 2^nd^_AtGDH-ΔT262-A263, and 7^th^_AtGDH-R246S crystals, respectively. BALBES, the automated (MR) pipeline, was used to identify the correct orientations of AtGDH molecules using the *C. symbiosum* GDH (CsGDH) (PDB ID: 1BGV) as a search model (https://ccp4online.ccp4.ac.uk) (Long et al., 2008). The latter shares 47.6% sequence identity with the WT AtGDH. BALBES run correctly placed two, six, two, and two template models in the correct orientation. Further, the AtGDH-I, AtGDH-II, 2^nd^_AtGDH-ΔT262-A263, and 7^th^_AtGDH-R246S structures were manually built in COOT (Emsley & Cowtan, 2004), and refinement cycles in REFMAC5 (Murshudov et al., 1997). Additionally, the amplitude and LORESTR pipeline (Kovalevskiy et al., 2016) refinement of the CCP4 suite was used occasionally.

Next, for the AtGDH-I, AtGDH-II, 2^nd^_AtGDH-ΔT262-A263, and 7^th^_AtGDH-R246S structures, the unresolved electron density peaks above 3σ level in the sigma-A weighted *F_o_-F_c_* electron density map were satisfied upon adding the solvent molecules. An iterative cycle of refinement and manual building in COOT was performed until all positive peaks in the electron density were satisfied and acceptable *R*_factor_ and *R*_free_ were obtained. The refinement statistics of these structures are presented in Table S1. The stereochemistry of the residues in all these built structures was analyzed by Ramachandran plot and PROCHECK (Laskowski et al., 1993). The PyMol Molecular Graphics System, Version 2.1.1, Schrödinger, LLC (DeLano, 2002), UCSF ChimeraX (Goddard et al., 2018), and UCSF Chimera 1.13 (Pettersen et al., 2004) were used for visualization, analysis, and figure preparation.

### Cryo-EM Sample Preparation and Data Collection for AtGDH and AnGDH samples

3µl of the AtGDH-AKG bound binary complex (3 mg/ml) was applied to freshly glow-discharged holey carbon grids (Quantifoil Au R 0.6/1.0 300 mesh). For the apo form AnGDH dataset, 6 mg/ml of AnGDH was mixed with 0.03% Tween-20 prior to application on the grid. The grids were incubated at 16 °C under 100% humidity and subsequently blotted for 4-4.5 sec. Immediately after blotting, the grids were cryo-cooled in liquid ethane on a Vitrobot Mark IV (Thermo Fisher Scientific). Before setting up data collection, the grids were carefully screened to assess the uniform particle distribution, any possible ice formation, and good quality density. The datasets were collected on a 300 kV FEI Titan Krios G3 transmission electron microscope equipped with the automated data collection software EPU (Thermo Fisher Scientific). The images of AnGDH and AtGDH-AKG binary complex were recorded at a defocus range of 2.1-3.3 µm on a Falcon III direct detector camera (ThermoFisher) operating in counting mode at 1.38 and 1.07 Å pixel^−1^, respectively. A total of 25 frames were collected with an electron dose of 27.67 and 25.75 e^-^/Å^2^ for each image of the AnGDH and AtGDH-AKG binary complex, respectively. The details of data collection parameters are given in the Table S2 below. Sample preparation and data collection were performed at the National Electron Cryo-Microscopy facility, Institute for Stem Cell Science and Regenerative Medicine (inStem), Bengaluru, India.

### Cryo-EM Data Processing

The image processing of the AnGDH and AtGDH-AKG binary complex datasets was performed in RELION3.1 and RELION4.0, respectively (Scheres, 2012). Initially, beam-induced motion of 1050 and 1902 images of AnGDH and AtGDH, respectively, were corrected using the MotionCor2 software program (Zheng et al., 2017). Further, the respective motion-corrected micrographs of AnGDH and AtGDH-AKG binary complex were subjected to the estimation of the Contrast Transfer Function (CTF) using the CTFFIND4 program (Rohou & Grigorieff, 2015). The micrographs were manually examined for the quality of CTF, thon rings, and particle distribution. Micrographs with aggregated protein were rejected. Initially, a small set of 20 micrographs were selected based on CTF values (low and high). Further, the parameters (box size, threshold values, etc.) were optimized for the auto-picking of particles. The particles were extracted from the selected template micrographs, and then 2D classification was performed. Further, the 2D class-averaged images in multiple orientations were selected and used as a template for reference-based auto-picking for both AnGDH and AtGDH-AKG binary complex data sets.

A 324964 particles were picked automatically from a total of 1025 micrographs. Further, the clustered particles and background were discarded. Finally, 282042 particles were selected from the 2D class averaging. Using the whole 2D class-averaged particles, the initial 3D map was generated in C1 symmetry. Further, the particles were 3D auto-refined, imposing D3 symmetry and using the 3D initial map as a reference. Further, 282042 particles were subjected to 3D classification with six classes. Out of six classes, two classes were rejected due to poorly reconstructed and conformational differences in the map. Further, the combined 3D classes containing 207704 selected particles were 3D auto-refined and post-processed (Figure S1).

For the AtGDH-AKG binary complex dataset, 398843 particles were picked automatically from 1902 micrographs. The clustered particles and background were discarded, and the remaining particles were subjected to 2D classification. Finally, single isolated 355523 particles were selected from the 2D class averaging. The initial 3D model of AnGDH was rescaled and used as a reference for 3D auto-refinement in C1 symmetry. Poor quality reconstructed maps were rejected. Further, the extracted 334277 particles were 3D auto-refined while imposing D3 symmetry. It was followed by 3D classification into four classes. The 3D class distribution was 15.4%, 27.9%, 24.3%, and 32.2%, with the 50757, 93686, 81516, and 108318 particles, respectively, assigned to each class. All the classes (1, 2, 3, and 4) were individually refined using the gold standard.

It was followed by masking and post-refinement of each refined map for both the AnGDH and AtGDH-AKG binary complex data sets. To improve the quality of the map, CTF refinement, and Bayesian Polishing were performed on both AnGDH and AtGDH-AKG binary complex data. The local resolution of the respective reconstructed maps of the AnGDH and AtGDH-AKG binary complex were calculated using the Local resolution tool in RELION3.0. All the generated maps were prepared and analyzed using the UCSF chimera or PyMol tool (Fig. S1, S2, and S3).

### Model Building for the cryo-EM Maps

The models of the AnGDH and AtGDH-AKG binary complex were manually built using the cryo-EM module of COOT. The crystal structure AnGDH (PDB ID: 5XVX) was used as a template to fit the AnGDH refined map. The crystal structures of AtGDH (AtGDH-II and PDB ID: 7ECS) were used as references for AtGDH-AKG binary complex structures. Further, the α-helices and β-strands were adjusted to the map density by the Jiggle fit chain, Chain Morph, or All Chain refine options in COOT (Emsley & Cowtan, 2004). Several iterations of real-space refinement in PHENIX (Afonine et al., 2012) further followed it.

After careful inspection of all the quaternary structures of the AtGDH-AKG bound binary complex, the density for the substrate AKG was found missing. Therefore, hereafter, the structure is considered as apo form.

### Molecular Dynamics (MD) simulation studies of AtGDH and AnGDH

All the molecular dynamics (MD) simulations in this study were performed by GROMACS 2022 (Van Der Spoel et al., 2005) with a charmm36 force field (Huang & MacKerell, 2013). AnGDH and AtGDH hexamers were individually placed at the center of the box at a distance of 10 Å from the wall, surrounded by SPC/E water molecules with periodic boundary conditions. Energy minimizations were performed for 50000 steps. The energy minimized systems were further equilibrated using canonical ensemble (NVT) followed by isothermal-isobaric ensemble (NPT). In the NVT equilibration, systems were heated to 300 K using V-rescale, a modified Berendsen thermostat for 1 ns. In NPT, all these heated systems were equilibrated using the Parrinello Rahman barostat for 1 ns to maintain a constant pressure of 1 bar. The unrestrained production MD simulations were performed for 300 ns for all structures. The covalent bonds involving H-atoms were constrained using the “LINCS” algorithm and the long-range electrostatic interactions with particle mesh Ewald (PME) method. The time step for integration was set to 2 fs during the MD simulation. GROMACS tools (Van Der Spoel et al., 2005), VMD (Humphrey et al., 1996), and Bio3D (Grant & Yao, 2021) were used for further analysis of the MD trajectory. Protein structure networks were obtained using Pyinteraph (Tiberti et al., 2014). The network graphs obtained were visualized using the open source software Cytoscape.

### Structural Analysis of AtGDH and AnGDH

The structures were analyzed using the COOT (Emsley & Cowtan, 2004) and visually inspected using PyMol software (DeLano, 2002). The open and partially closed conformations of AtGDH-I (subunits - A and B) and AnGDH (5XVI, subunits - A and E) were compared. The Least-Square (LSQ) Fit Superposition was done for the regions: Domain I (1-190), Domain II (192-375), 375-400, hinge helix (400-436), and C terminus of Domain I (437-460) between the open to partially closed conformations of AtGDH and AnGDH. The Domain I of the open and partially closed conformation were superposed to identify the differential Domain II motion between AnGDH and AtGDH. The Domain II motions with respect to its rotation about the hinge helix α15 and the α7 helix movement were critically analyzed on transition from an open to a partially closed state. For the α7 helix movement, the degree of rotation was calculated between the conserved residues A196 and G215 (open sate) to G215 (partially closed state).

## Results

### Quality of the crystal and cryo-EM structures of AtGDH and AnGDH and their overall structural fold

We have determined multiple structures of AtGDH using X-ray crystallography and cryo-EM. The AtGDH crystals were obtained under two different conditions, leading to the determination of the apo forms of AtGDH crystal structures (AtGDH-I and AtGDH-II) at resolutions of 2.0 and 2.85 Å, respectively. Both AtGDH-I and AtGDH-II structures have been successfully refined, as indicated by the acceptable *R*_factor_ and *R*_free_ (Table S1). We have also determined crystal structures of AtGDH mutants - 2^nd^_AtGDH-ΔT262-A263 and 7^th^_AtGDH-R246S, refined with reasonable final *R*_factor_ and *R*_free_ values (Table S1).

Additionally, we report the first cryo-EM structures of the fungal GDHs from *A*. *niger* (Fig. S1) and *A. terreus* (Fig. 1A, 1B, S2 and S3) (Table S2). The cryo-EM structure of apo form AnGDH was determined. The AnGDH reconstructed map resolution was predicted by the gold standard (0.143 cut-off) Fourier Shell Correlation (FSC) to be 3.6 Å (Fig. S1E). The final correlation coefficient (CC) for AnGDH is 0.7, which indicates an acceptable model (Table S2). Similarly, four cryo-EM structures of AtGDH were determined. The resolutions of 3-dimensional (3D) reconstructed maps for AtGDH ensembles were (FSC 0.143 cut-off) were 3.65, 3.15, 3.25, and 3.18 Å, respectively (Fig. S2 and S3). The reconstructed cryo-EM maps of AtGDH displayed four distinct quaternary structural conformations (referred to as AtGDH-II-em1, AtGDH-II-em2, AtGDH-III-em1, and AtGDH-III-em2) (Fig. S2) (Movie 1). The final correlation coefficient (CC) for AtGDH-II-em1, AtGDH-II-em2, AtGDH-III-em1, and AtGDH-III-em2 were 0.8, 0.54, 0.8, 0.7, and 0.72, respectively, thus indicating good quality models (Fig. 1B) (Table S2).

**Figure 1.**
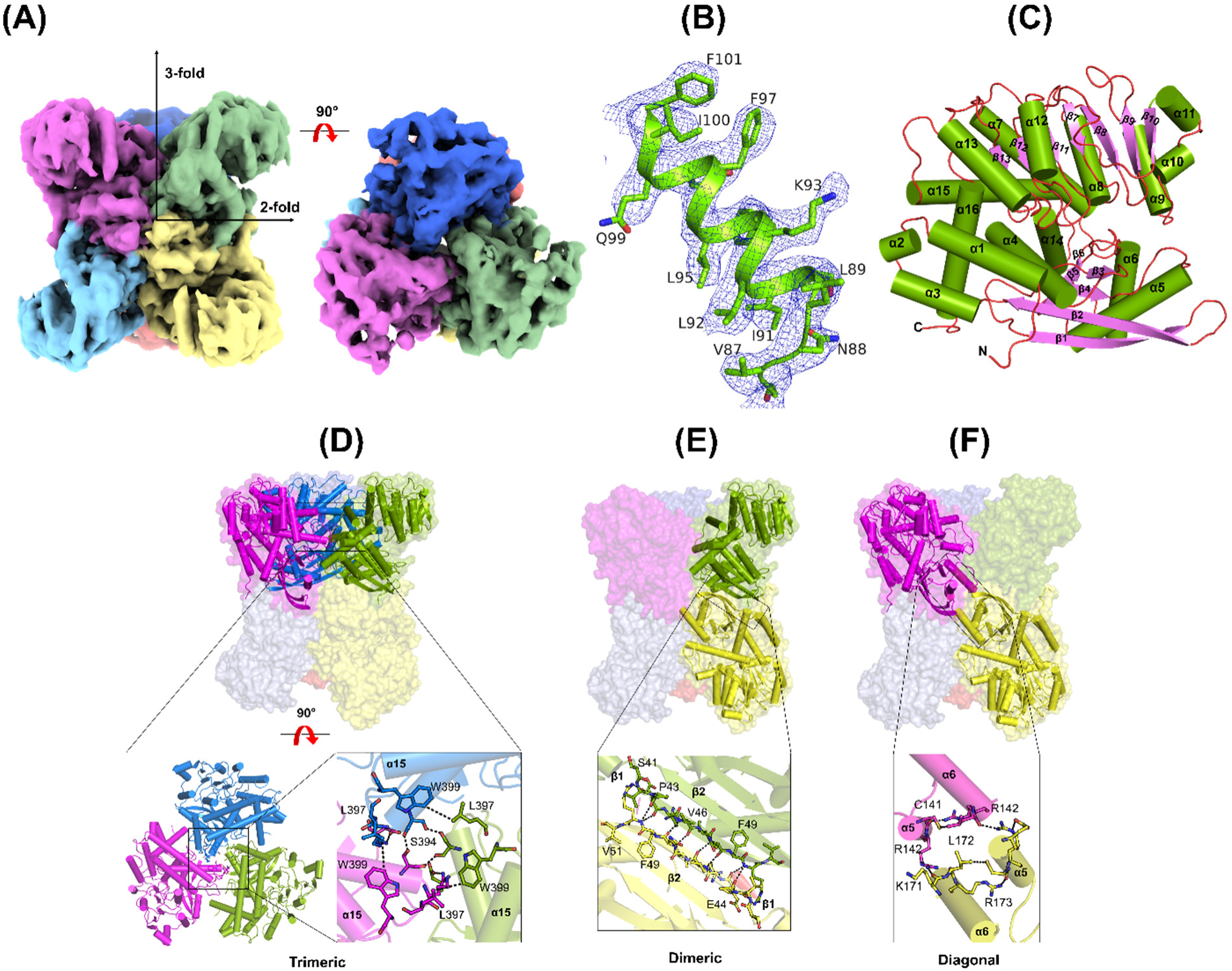
Cryo-EM structure of AtGDH and its inter-subunit interactions. (**A**) Cryo-EM map (different colors for different subunits) of hexameric AtGDH reconstructed with applied D3 symmetry. A 90° rotated view of the map displays the 3-fold symmetry arrangement of the hexamer. (**B**) Real space electron density at > 5σ contour level is shown as blue mesh around the α helix along with the side chains of AtGDH model. (**C**) Cartoon representation of one subunit displays the tertiary structural fold of AtGDH. The α-helices (lime green) are labeled as ‘α’, β-strands (purple) as ‘β’, with their numbers. The loops are shown in brown color. Representative views of interfacial interactions along the trimeric (**D**), dimeric (**E**), and diagonal axis (**F**) are shown for a hexamer. The interacting subunits are displayed as cylindrical cartoons in a hexamer (transparent surface). Insets display the interacting residues shown as sticks at the trimeric, dimeric or diagonal interfacial regions.

The overall structural folds of the cryo-EM structures of AnGDH and AtGDH are similar to the previously reported crystal structures of the same fungal enzymes (Prakash et al., 2018; Godsora et al., 2022) and hexameric GDHs from other organisms (Baker et al., 1992; Peterson & Smith, 1999; Grzechowiak et al., 2020) (Fig. 1C). Each subunit of AtGDH and AnGDH has two domains (I and II). The hexameric AtGDH and AnGDH (Fig. 1A) are a ‘dimer of trimers.’ Several inter-dimeric and trimeric interactions form the functional hexameric unit. The inter-trimeric interactions are formed by β_3_, β_5_, β_6_, α_14_, and α_16_ (Fig. 1D), while the inter-dimeric interactions are formed by β_1_, β_2_, and α_5_ (Fig. 1E). Additionally, the diagonal interactions are formed between the top and bottom subunits *via* two helices (α5 and α6) from both subunits arranged in a diagonal manner (Fig. 1F).

### Conformational heterogeneity of AtGDH and AnGDH

Intrinsic motion in the enzymes is associated with their functional properties. We have captured one conformation of AnGDH and three distinct conformational states of AtGDH (AtGDH-I, II, and III), as presented in Figure 2. Based on the opening of the active site cleft (mouth opening), which is measured by the distance between the Cα atoms of the residues - K122 and R282 (R280 in AnGDH), the acquired conformations have been categorized as open (17-21 Å), partially closed (10-15 Å), or closed (7-8 Å) states (Table S3). The cryo-EM structure of AnGDH has all the subunits in a closed conformation (average mouth opening, ∼8 Å). This is the first structure of AnGDH to be reported in a closed conformation, even without any substrate.

In the crystal forms, we have captured two distinct quaternary structures of AtGDH: (a) AtGDH-I having subunits in open (average mouth opening, ∼17 Å) and partially closed (average mouth opening, ∼13 Å) conformations (Fig. 2A), and (b) AtGDH-II with all subunits in partially closed conformations (average mouth opening, ∼12 Å) (Fig. 2B). Similarly, in the solution, two cryo-EM structures (AtGDH-II-em1 and AtGDH-II-em2) of AtGDH also exhibit partially closed conformations (average mouth opening, ∼12 Å) (Fig. 2B and 2D). Another predominant conformational ensemble of AtGDH observed in cryo-EM structural data has all the subunits in the closed conformations (average mouth opening, ∼8 Å), designated as AtGDH-III-em1 and AtGDH-III-em2 (Fig. 2C). In total, we have determined six structures (including two crystal structures and four cryo-EM structures), however, on the basis of their structural features, they belong to either of these three quaternary structural forms, and a systematic analysis of each structural assembly is presented below.

**Figure 2.**
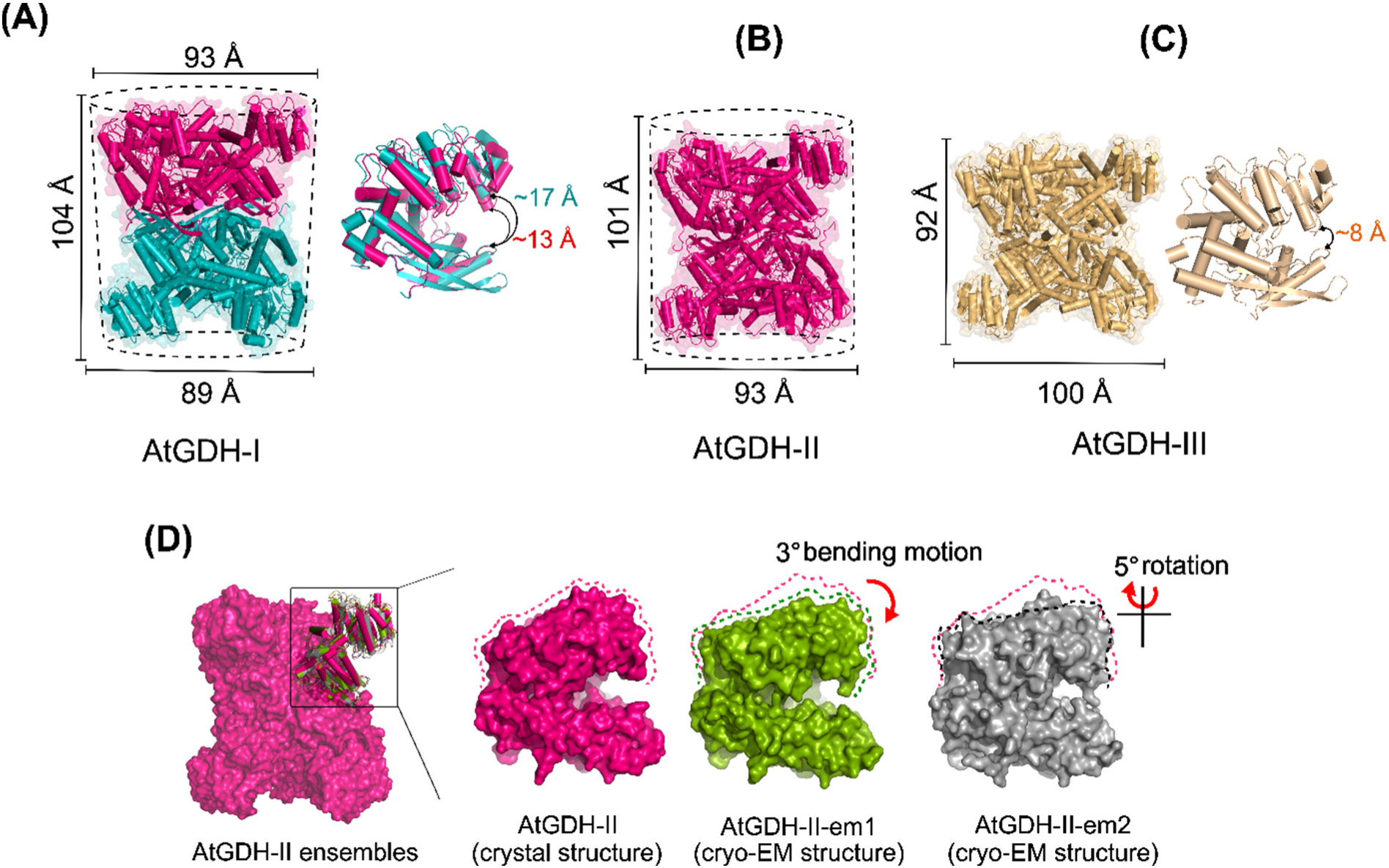
Conformational dynamics of the subunits in hexameric apo- AtGDH crystal and cryo-EM structures. Cartoon and surface (transparent) representations of the hexameric AtGDH. The dimensions of the hexameric functional assembly are labeled in terms of height (left) and diameter (top and bottom). The dashed lines around the structures (panels A and B) depict their visual shape. (**A**) AtGDH-I crystal structure has two co-existing conformations, partially closed (dark pink, ∼13 Å) and open conformation (teal, ∼17 Å), stacked on each other. Conformational differences in the monomers are also depicted on the side. (**B**) AtGDH-II structures (AtGDH-II crystal, AtGDH-II-em1, and AtGDH-II-em1 cryo-EM) have all the subunits in the partially closed conformation (dark pink, ∼11-13 Å). (**C**) The ensembles of AtGDH-III structure (AtGDH-III-em1 and em2 cryo-EM) have all the subunits in closed conformations (light orange) with an average mouth opening of ∼7-8 Å. The mouth opening distance of one subunit (cartoon) is shown. (**D**) Surface representation of AtGDH-II ensemble structures. The cartoon representation depicts superposed subunits of AtGDH-II ensembles. One subunit of each ensemble: AtGDH-II crystal structure (dark pink) and cryo-EM structures - AtGDH-II-em1 (green) and AtGDH-II-em2 (gray) are shown as surface. The dashed lines (pink, green, and black) highlight the differences in Domain II movement due to bending and shear motions. The observed bending and shear motions in the subunits of AtGDH-II-em1 and AtGDH-II-em2 ensembles are shown with respect to the AtGDH-II crystal structure (pink dashed line).

Hexameric AtGDH can be represented as a ‘cylinder’. AtGDH-I structure with subunits having both open and partially closed conformations appear like distorted cylinder, with an average height of 104 Å, and diameter of 89 Å at one side (open state trimer) and 93 Å at the other side (partially closed trimer) (Fig. 2A). The ensembles of AtGDH-II structure (AtGDH-II crystal and cryo-EM: AtGDH-II-em1, and AtGDH-II-em2) have all the subunits in partially closed conformation, and have an appearance like a perfect cylinder (Fig. 2B) (Table S3). The calculated r.m.s.d. values of the superposition of AtGDH-II crystal to AtGDH-II-em1 and AtGDH-II-em2 are 2.3 and 1.3 Å, respectively. Mostly, the active site cleft opening for the subunits in these hexamers ranges from ∼10-13 Å, while in AtGH-II-em1, one of the subunits adopts a mouth opening of ∼15 Å (Table S3). Despite having the exact mouth opening distances between AtGDH-II structural ensembles, slight variations in Domain II movement are observed that are referred to as ‘bending’ and ‘rotation’ motions (Fig. 2D). These AtGDH-II ensemble structures have an average height of 101 Å and a diameter of 93 Å (Fig. 2B).

Furthermore, other structural forms of AtGDH, i.e., AtGDH-III-em1 and AtGDH-III-em2, both have all the subunits adopting only closed (∼7-8 Å) conformation (Fig. 2C). It forms a horizontally expanded quaternary structure (perfect cylinder appearance) with an average height and diameter of 92 and 100 Å, respectively (Fig. 2C). Like AtGDH-II ensembles, the AtGDH-III ensembles structure with the exact mouth opening distances, display bending and rotation motion in Domain II. Notably, the helix α7 moves upward by 5.9 Å (reference point T197), and the hinge helix (α15) slides away from the trimeric center by 3.6 Å (reference point K424), leading to the shear motion of Domain II. Thus, the captured multiple quaternary structures of apo-AtGDH and apo-AnGDH (including the previously reported structures by Prakash et al., 2018) demonstrate conformational heterogeneity in both these enzymes in apo state. Further, the comparative studies between these enzymes will help to justify the kinetic discrepancies.

### Modulation of the structural elements, surface charges, and interacting networks in Domain II due to the amino acid replacements between wild type AtGDH and AnGDH

Comparative studies between wild types (WT) AtGDH and AnGDH (share 88 % amino acid sequence identity shown in Figure S4) were performed. The inter-subunit interactions between AtGDH and AnGDH along the inter-dimeric and -trimeric axis are conserved. However, few substitutions in helices α5 (residues - S127N, A135S, and R142K), α6 (residues - I162V, Y164F, and L172I), and the connecting loop (residue S175Q), which participate in the diagonal interactions are present in AtGDH.

Notably, most of the amino acid substitutions (10 %) reside in Domain II and hinge helix (Fig. 3A and 3B), which undergoes significant conformational changes during the opening and closing of the active site cleft for catalysis. Few amino acid substitutions have caused alterations of the surface charge distribution as well as the hydrogen bonding network between AnGDH and AtGDH, particularly at a few specific regions, i.e., insertion (near Loop 1) and deletion (Loop 2) found in AtGDH (Fig. 3C, 3D, and S5). In AnGDH, the conserved R223 lies closer to Loop 1 and interacts with E262. However, to accommodate two amino acid insertions (T262 and A263 in Loop 1) in AtGDH, the conserved R223 flips 120° outward. The re-oriented conserved R223 resides closer to the R246 residue, thereby creating a positively charged groove on the AtGDH surface (Fig. 3C). In contrast, the same region has a neutral surface charge distribution on AnGDH, as S246 substitutes R246 (Fig. 3D).

**Figure 3.**
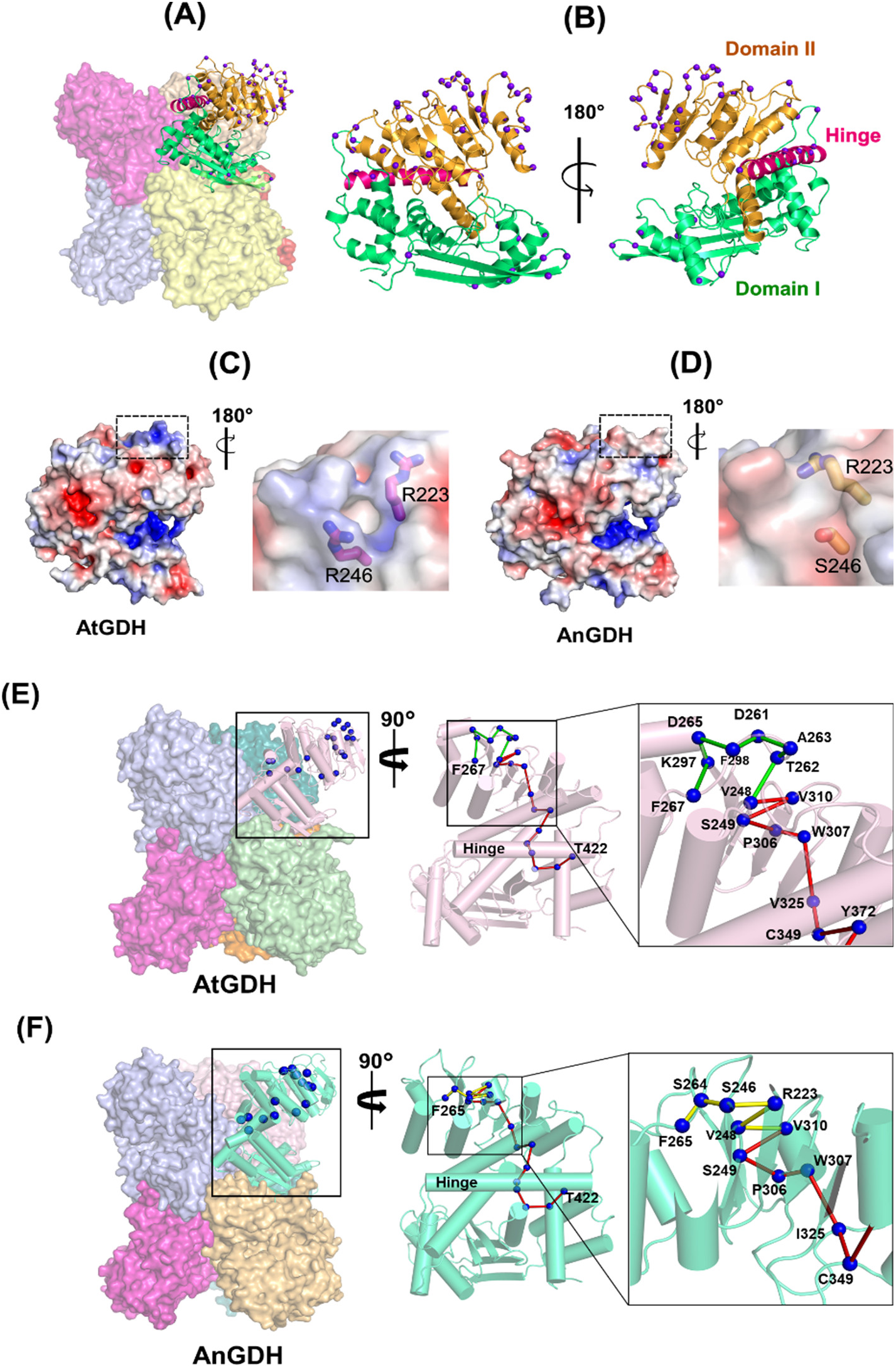
Impact of amino acid substitutions on the surface charge distribution and intrinsic network. **(A)** One subunit hexameric AtGDH (surface) is shown as a cartoon, and the locations of the amino acid substitutions as compared to AnGDH are shown as a sphere. **(B)** The front and rear views of the monomer are shown. The purple-blue sphere represents the positions of the substituted amino acids between AtGDH and AnGDH. Electrostatic charge distribution on the overall surface of AtGDH **(C)** and AnGDH **(D)** monomers are shown. The dashed box marks the notable differences in the charge distribution at the same position on AtGDH and AnGDH. Altered orientation of the conserved R223 residue in AtGDH (**C**, inset) and AnGDH (**D**, inset) and its impact on the surface charge distribution. The red colored surface indicates negative charge, and the blue colored surface indicates positive charge. A proposed network connecting the hinge helix to the edge of Domain II in one subunit of AtGDH (colored light pink) (**E**) and AnGDH (colored cyan) (**F**). The residues are represented as blue spheres with red, green, and yellow bonds connecting them. The conserved network pathway between AnGDH and AtGDH is connected by red bonds. The non-conserved pathway between AtGDH and AnGDH, is connected by green bonds (in AtGDH) and yellow bonds (in AGDH). The zoomed-in view of the non-conserved pathway is shown, and the residues are numbered.

The tip of Domain II is guarded by residues ranging from S266-R305, which account for the highest number of amino acid substitutions between AtGDH and AnGDH. It is the most non-conserved region in GDHs characterized in various living organisms. Therefore, we hypothesized that a long-range well connected network might exist between the tip of the cleft (α9) and the hinge helix (α15). Several interacting network routes might co-exist to perform the Domain II associated opening and closure during the catalysis.

The calculated r.m.s.d and root mean square fluctuation (r.m.s.f) analysis from the MD simulations run indicated AtGDH to be a more dynamic enzyme than AnGDH (Fig. S6A and S6B). Based on dynamic protein structure network (PSN) analysis performed using Pyinteraph (Tiberti et al., 2014), we identified unique interactions within a subunit of AnGDH and AtGDH. Notably, two interactions, D265-K297 and D265-D261, were present only in AtGDH and not in AnGDH. Our analysis suggests a network connecting the hinge helix with the edge of Domain II involving unique interactions. For AtGDH the network is: T422-A444-L105-C415-A377-P374-Y372-C349-V325-W307-P306-S249-V310-V248-T262-A263-D261-D265-K297-F298-F267 (Fig. 3E). For AnGDH the network is: T422-A444-L105-C415-A377-P374-Y372-C349-I325-W307-P306-S249-V310-V248-R223-S246-S264-F265 (Fig. 3F). Comparison of the networks connecting the residues T422 to F265 in AnGDH and F267 in AtGDH displays a conserved pathway from T422 to V248 (colored red in Fig. 3E and 3F). The pathways from V248 to F265 in AnGDH and F267 in AtGDH are different (colored yellow in AnGDH and green in AtGDH) (Fig. 3E and 3F). This is because of the presence of the extended loop (Loop 1) in AtGDH consisting of the residues T262-A263 in close proximity. In addition to this, K297, located on the small helix in AtGDH next to the extended loop, is a part of the network, unlike in AnGDH. In addition to these, the interaction S246-S264 was observed predominantly in AnGDH and not in AtGDH (Weight of the interaction: AnGDH = 99%; AtGDH = 12%). Therefore, the network pathway connecting T422 to F265 or F267 revealed a few crucial residues. For example, the insertion in AtGDH consists of residues D261-D265 (Loop 1) and, K297, and S246 in AnGDH. Together, the comparative structural analysis between allosteric AnGDH and non-allosteric AtGDH highlighted that the substitutions on Domain II modifies the electrostatic charge distributions and the interacting network extending from the hinge helix to tip region. So, some of these substituted residues were thus further considered for site-directed mutagenesis to confirm their involvement in the kinetic regulations of these GDHs.

### Variance in Domain II dynamics between AtGDH and AnGDH controls the transition from open to partially closed conformation

Impact of the altered network in Domain II on the conformational dynamics and inter-subunit interactions was deduced. Systematic analysis of the structural changes related to the transition process from the open to partially closed conformations was performed for both apo forms of AnGDH (PDB ID: 5XVI, Prakash et al., 2018) (Fig. 4A and 4B) and AtGDH (AtGDH-I) (Fig. 4E and 4F). A significant disparity in the extent of mouth opening was observed in the open conformations of AtGDH and AnGDH, while the closed and partially closed conformations were alike. Notably, there is a difference of ∼4 Å in the active site cleft opening between the open conformations of AnGDH (∼ 21 Å) and AtGDH (∼17 Å) (Fig. 4B and 4F).

**Figure 4.**
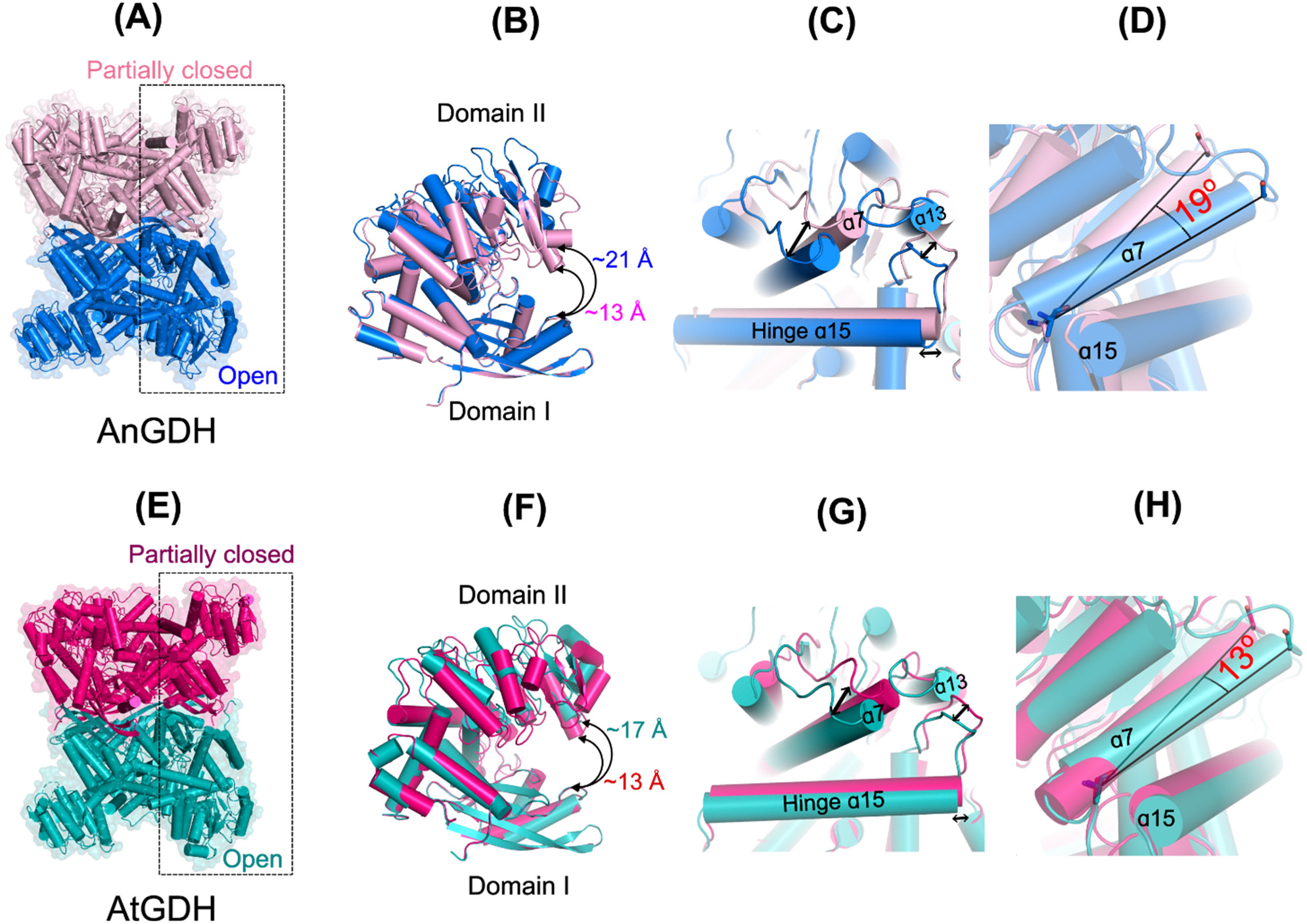
Domain II motion during open to partially closed conformational state transition in AnGDH and AtGDH. The rotation of helix α7 is obtained by measuring the angle between the residues A196 and G215 (from the open state) to G215 (partially closed state). The residues A196 and G215 are shown as sticks with respective color codes. (**A**) Hexameric AnGDH (PDB ID: 5XVI) with co-existing partially closed (light pink) and open conformations (blue). (**B**) Superposition of AnGDH subunits (subunit A and subunit F marked in dashed box) (PDB ID: 5XVI) existing as partially closed (light pink) to open conformations (blue). (**C**) The double-headed arrow indicates the movement of the hinge helix α15 and neighboring loops during the structural transition from the open to the partially closed conformation. (**D**) The 19° rotation of α7 helix during the transition from an open to a partially closed conformation in AnGDH. (**E**) AtGDH-I has two co-existing conformations – partially closed (dark pink) and open conformations (teal). (**F**) Superposition of AtGDH subunits (subunit A and subunit B) marked in dashed box existing in partially closed (pink) to open conformations (teal). (**G**) The double-headed arrow symbol shows the movement of the hinge helix α15 and the neighboring loops. (**H**) The 13° rotation of α7 helix during the transition in AtGDH.

In allosteric AnGDH, when the monomer switches from open (mouth opening ∼21 Å) to partially closed (mouth opening ∼13 Å) conformation (Fig. 4A and 4B), the ascending α7 helix of Domain II shifts by ∼19° with respect to the hinge helix (α15) (Fig. 4D). Even the α14 helix, which forms part of the active site, undergoes a significant conformational change. Further, the flexible hinge helix (α15) slides by 2 Å away from the trimeric center (Fig. 4C). Furthermore, the connecting loop (Y426-S436) extending from the back of the hinge helix (α15) moves forward by 6.0 Å (Fig. 4C). On the contrary, the secondary structural elements of non-allosteric AtGDH exhibit relatively minor changes, between the open (∼17 Å) and partially closed (∼13 Å) conformations (Fig. 4E and 4F). The ascending α7 helix of Domain II in AtGDH shifts by ∼13° (Fig. 4H). The flexible hinge helix (α15) shifts slightly away from the trimeric center by 1.7 Å. Moreover, the connecting loop (426–436) extending from the hinge helix moves forward by 5.5 Å (Fig. 4F). Additionally, the dynamicity was localized substantially to the Domain II residues in both enzymes (Fig. S6A and S6B). AnGDH displays rigid body motion of the Domain II along the hinge helix. On the contrary, AtGDH displays several side chain flipping of residues in Domain II in addition to hinge movement (Movie S2 and S3).

The differential conformational dynamics affect the inter-subunit interactions between the open and partially closed subunits of AtGDH and AnGDH. The inter-subunit interactions between conserved residues - K171 and Q456 exist only in AnGDH. Moreover, the S175-D147 inter-subunit interaction is unique to AtGDH (Table S4). Thus, the structural analysis presented in the above sections indicates that residue substitutions in Domain II affect the conformational dynamics, leading to altered interactions and compactness at the inter-subunit regions of allosteric AnGDH and non-allosteric AtGDH (Fig. S7). It is also evident that the amino acid substitutions located in the Domain II and hinge helix regions might be crucial for regulating the dynamicity of these two enzymes. Mutational studies and kinetic characterizations of the mutant enzymes were performed to probe into the role of these substitutions towards different kinetic properties of AnGDH and AtGDH.

### AtGDH and AnGDH mutants exhibit altered kinetic properties

Site-directed mutagenesis in AtGDH and AnGDH were performed. The residues for mutagenesis were selected based on either of these criteria, such as closeness to the hinge helix α15, role in exhibiting surface charge differences, substitutions in the inter-subunit region, insertion or deletion regions, or involvement in salt bridge formation or from the differences observed in the dynamic network analysis within the subunits.

All the AtGDH and AnGDH mutants were successfully expressed and purified as a hexamer (MW ∼300 kDa) (Fig. S8). All the AtGDH and AnGDH mutants showed no changes in the secondary structural elements (Fig. S9) compared to the corresponding wild type enzymes. Eight sequential site-directed mutagenesis were performed on AtGDH to make this enzyme kinetically cooperative with the substrate AKG (Fig. S10). Mutations were considered based on the substitutions present in AnGDH. Initially, the amino acid insertion sites (T262-A263) (Fig. S5) were selected, followed by the neighboring residues contributing to the differential surface charge (Fig. 3C and 5A). The deletion mutant (2^nd^_AtGDH_ΔT262-A263) and the sixth mutant (6^th^_AtGDH-KDTAKD) showed slightly lower enzyme activity than the WT AtGDH (Fig. 5A and 5B) (Table 1). Notably, the sixth mutant lost the substrate inhibition property of WT AtGDH (Table 1).

**Figure 5.**
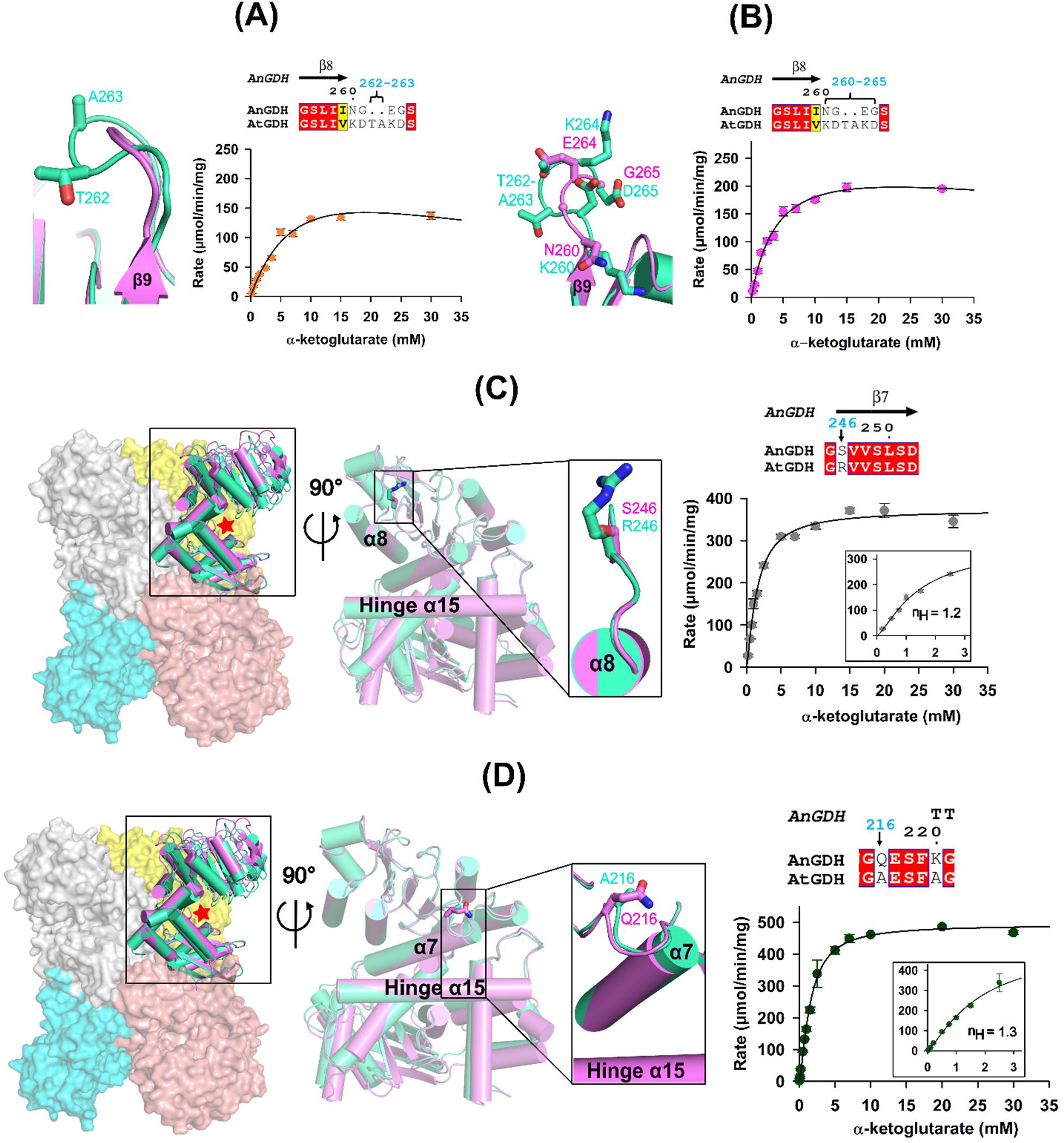
Role of amino acid insertion and charged substitutions in the kinetic behavior of AtGDH. In each panel, the amino acid sequence alignment of AtGDH and AnGDH on top displays the point of mutations (marked with a black arrow and residue number written in the blue label) performed in AtGDH. The red box indicates the conserved regions. Surface representation of hexameric AtGDH displaying the locations of these mutations is shown in Figure S10. Superposed structure of WT AtGDH (lime green carbon) and WT AnGDH (magenta carbon) are shown on left side of each panel. The substituted residues are shown in ball and stick in their respective color code. (**A**) The zoomed inset represents the orientation of Loop 1, which has two amino acid insertions- T262 and A263 in AtGDH as compared to AnGDH. Kinetic plot representing the AKG substrate saturation of the 2^nd^_AtGDH-ΔT262-A263 mutant (right side). **(B)** The zoomed inset represents the substituted charged residues (K262, D261, K264, and D265) in AtGDH as compared to AnGDH. Kinetic plot representing the AKG substrate saturation of the 6^th^_AtGDH- KDTAKD mutant shown in a magenta circle. Surface representation of hexameric AnGDH (shown in different colors) in panels C and D. Inset shows the 90-rotated view of mutation location in both panels **C** and **D**. Kinetic characterization of 7^th^ _AtGDH-R246S (**C**) and 8^th^_AtGDH-A216Q (**D**) mutants are shown. Insets (panels C and D) display the zoomed-in view of the reaction velocities at the lower AKG concentration. The data points are the calculated initial velocities at the respective substrate concentration used, which was considered for all the AtGDH mutants. The calculated initial velocities for the AtGDH mutants were performed against the same substrate concentrations. The standard error is calculated for the three data sets shown. The curve represents the best fit in the non-linear regression plot in SigmaPlot. The correlation coefficient (R^2^) is greater than 0.95.

**Table 1.** α-ketoglutarate saturation of AtGDH and AnGDH mutants.

| AtGDH |  |  |  |  |  |  |  |
| --- | --- | --- | --- | --- | --- | --- | --- |
| | No. of mutations | $V_{\max}$<br>($\mu\text{mol}/\text{min}/\text{mg}$) | $S_{0.5}$ or $K_m$<br>(mM) | $n_H$ | $K_i$<br>(mM) | $k_{\text{cat}}$ ($\text{sec}^{-1}$) | $k_{\text{cat}}/K_m$<br>($\text{sec}^{-1} \text{M}^{-1}$) |
| WT | - | $329.3 \pm 100$ | $4.3 \pm 1.5$ | - | $45.8 \pm 8$ | 1646.5 | $3.8 \times 10^5$ |
| $\Delta\text{T262-A263}$ | 2 | $294.8 \pm 54.5$ | $10.2 \pm 2.7$ | - | $35.8 \pm 15.2$ | 1474.0 | $1.4 \times 10^5$ |
| K260N-D261G-K264E-D265G | 6 | $272.9 \pm 22$ | $4.3 \pm 0.7$ | - | $120 \pm 55$ | 1364.5 | $3.2 \times 10^5$ |
| R246S | 7 | $445.7 \pm 21$ | $2.3 \pm 0.24$ | 1.2 | $191 \pm 83$ | 2228.5 | $9.7 \times 10^5$ |
| A216Q | 8 | $663.1 \pm 33$ | $2.8 \pm 0.3$ | 1.3 | $89.3 \pm 22$ | 3315.5 | $1.2 \times 10^6$ |
| AnGDH |  |  |  |  |  |  |  |
| WT* | - | $110 \pm 10$ | $5.6 \pm 0.3$ | 2.9 | - | 550.0 | $0.9 \times 10^5$ |
| S246R | 1 | $82.7 \pm 2.9$ | $10.2 \pm 0.5$ | $2.1 \pm 0.12$ | - | 413.5 | $0.4 \times 10^5$ |
| Q216A | 1 | $79.7 \pm 4.08$ | $10 \pm 0.63$ | $2.7 \pm 0.31$ | - | 398.5 | $0.4 \times 10^5$ |

\* The experimental values are obtained from Agarwal et al., 2019.

The following sequential mutations were done at positions-R246S and A216Q, which reside on the neighboring loops in correspondence to the previous mutations (Fig. 5C, 5D, and S10). These loops are highly dynamic and undergo a conformational change during the active site cleft opening and closing (Movie S2 and S3). These serial AtGDH mutants – 7^th^_AtGDH-R246S (Fig. 5C) and 8^th^_AtGDH-A216Q (Fig. 5D) and have individually increased the catalytic efficiency by 2- to 3- fold (Table 1). Interestingly, 7^th^_AtGDH-R246S and 8^th^_AtGDH-A216Q mutants showed a gain in cooperative behavior towards the substrate AKG (Fig. 5C and 5D). Thus, these sequential mutations of residues residing far away from the active site have affected the *V*_max_, *K*_m_, or *k*_cat_/*K*_m_ of the non-allosteric AtGDH enzyme (Table 1). Additionally, two mutations (R246S and A216Q) converted the AtGDH enzyme to an allosteric one.

We thus decided to examine the two substitutions (R246S and A216Q), which drastically affected the kinetic properties of AtGDH on the allosteric AnGDH. Two AnGDH mutants (AnGDH-Q216A and AnGDH-S246R) individually showed 2-fold lower catalytic efficiency compared to WT AnGDH (Table 1) (Fig. 6A and 6B). However, it is remarkable to observe that the Hill coefficient values of AnGDH-S246R (*n*H = 2.1) and AnGDH-Q216A (*n*H = 2.7) mutants were lower than the WT AnGDH (*n*H = 2.9) (Fig. 6) (Table 1). These data indicate that AnGDH-S246R and AnGDH-Q216A mutants have lost some level of AKG dependent cooperativity due to these mutations (Table 1). Thus, these observations imply that the residue S246 must be playing a crucial role in allosteric transmission for structural modulation required for AKG binding in WT AnGDH and AtGDH mutant. So, structural studies of the AtGDH mutants will provide more insights into the network responsible for the differences in the kinetic properties of AtGDH and AnGDH.

**Figure 6.**
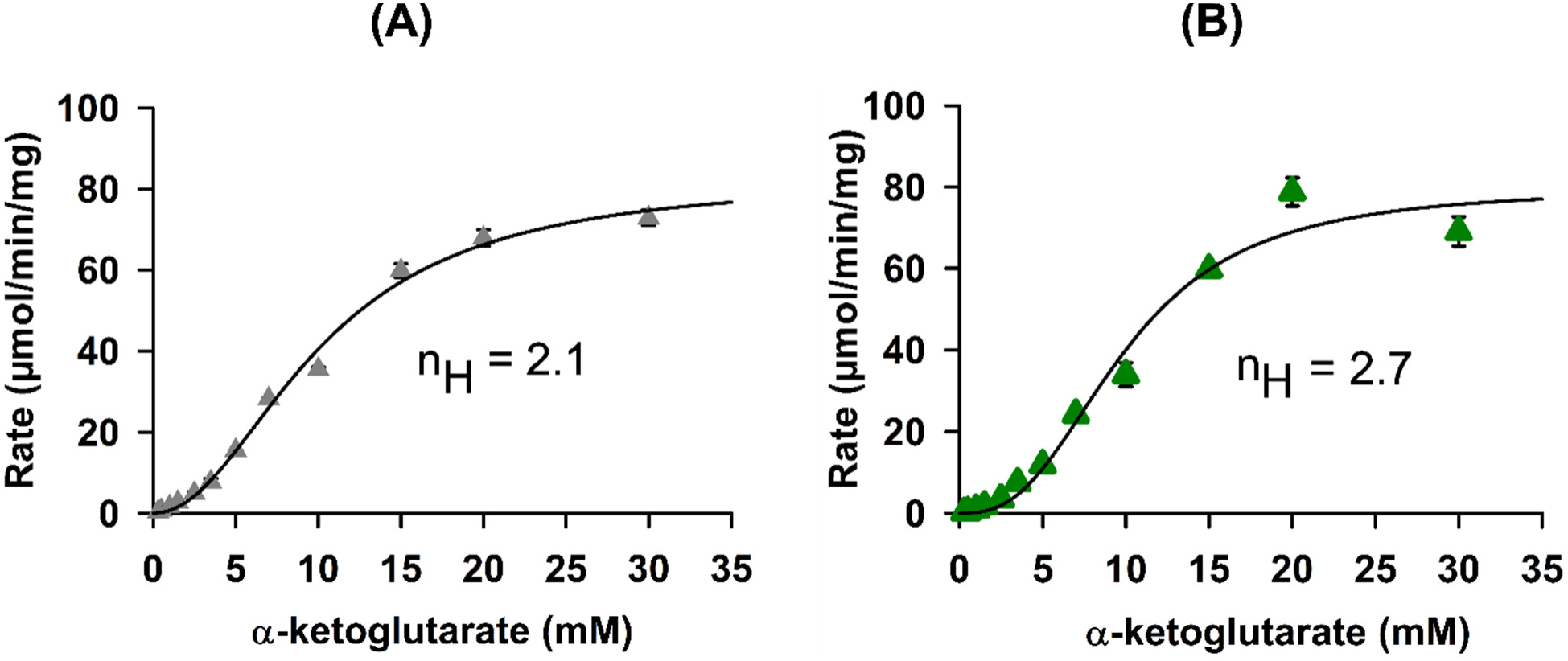
α-ketoglutarate saturation of AnGDH mutants and their fit to Hill equation. Kinetic characterization of AnGDH-S246R mutant (**A**) and AnGDH-Q216A mutant (**B**). The data points (shown as gray or green triangles) are the calculated initial velocities at the respective substrate concentration used. The standard errors are calculated for the three data sets shown. The curve represents the best fit in the non-linear regression plot in SigmaPlot. The correlation coefficient (R^2^) is greater than 0.95.

### Evidence of an altered network as observed in the crystal structure of the AtGDH mutants

The crystal structures of the 2^nd^_AtGDH-ΔT262-A263 and 7^th^_AtGDH-R246S mutants were solved (Table S1). Both the AtGDH mutants have 458 amino acids instead of 460 amino acids. The 2^nd^_AtGDH-ΔT262-A263 structure has two amino acid deletions (T262 and A263) (Fig. S11). While the 7^th^_AtGDH-R246S mutant has seven mutations at positions - R246S and Loop 1 (consisting of K260N, D261G, ΔT262-A263, K264E, D265G) in correspondence to substitutions found in AnGDH (Fig. S10 and S12).

Like WT AtGDH-I, the functional hexameric units of both the mutants have two conformational sub-states in the asymmetric unit. The mouth openings of the open and partially closed states in the 2^nd^_AtGDH-ΔT262-A263 structure are ∼14.4 and 17.2 Å, respectively (Fig. S11A), while ∼15.7 and 19.3 Å, respectively, in the 7^th^_AtGDH-R246S mutant (Fig. S12A). Comparative analysis of open conformations of AtGDH mutant structure with the WT AtGDH and WT AnGDH were performed (Fig. 7A). The overall structural fold is identical to the WT AtGDH except at the Loop 1 and Loop 2 regions (Fig. 7B and 7C). In the 2^nd^_AtGDH-ΔT262-A263 structure, the shortened mutated Loop 1 (ΔT262-A263) acquires two alternate conformations, and one coincidently matches the WT AnGDH (Fig. S11B and S11C). The neighboring residues - R246 and R223, rearrange in the vicinity in accordance with the conformation of mutated (ΔT262-A263) Loop 1 in the 2^nd^_AtGDH-ΔT262-A263 structure (Fig. S11C). On the contrary, the mutated Loop 1 (260-265) of the 7^th^_AtGDH-R246S structure adopts conformation similar to WT AnGDH (Fig. S12 and 7C). The conserved R223 in 7^th^_AtGDH-R246S mutant (Fig. 7B, 7D and 7E, blue carbon) has an orientation identical to WT AnGDH (Fig. 7E, light pink carbon). The mutated E262 (K264 to E) occupies the same position as found in WT AnGDH (Fig. 7E) and interacts with the conserved R223. Mutations on Loop 1 induce structural fold differences as well as consecutive interactions with Loop 2 in the 7^th^_AtGDH-R246S mutant (Fig. 7C, 7F, and 7G); however, no observed changes in the 2^nd^_AtGDH-ΔT262-A263 structure. The 7^th^_AtGDH-R246S mutant exhibits an altered network in Domain II, which is a combination of AtGDH and AnGDH networks (Fig. 7H).

**Figure 7.**
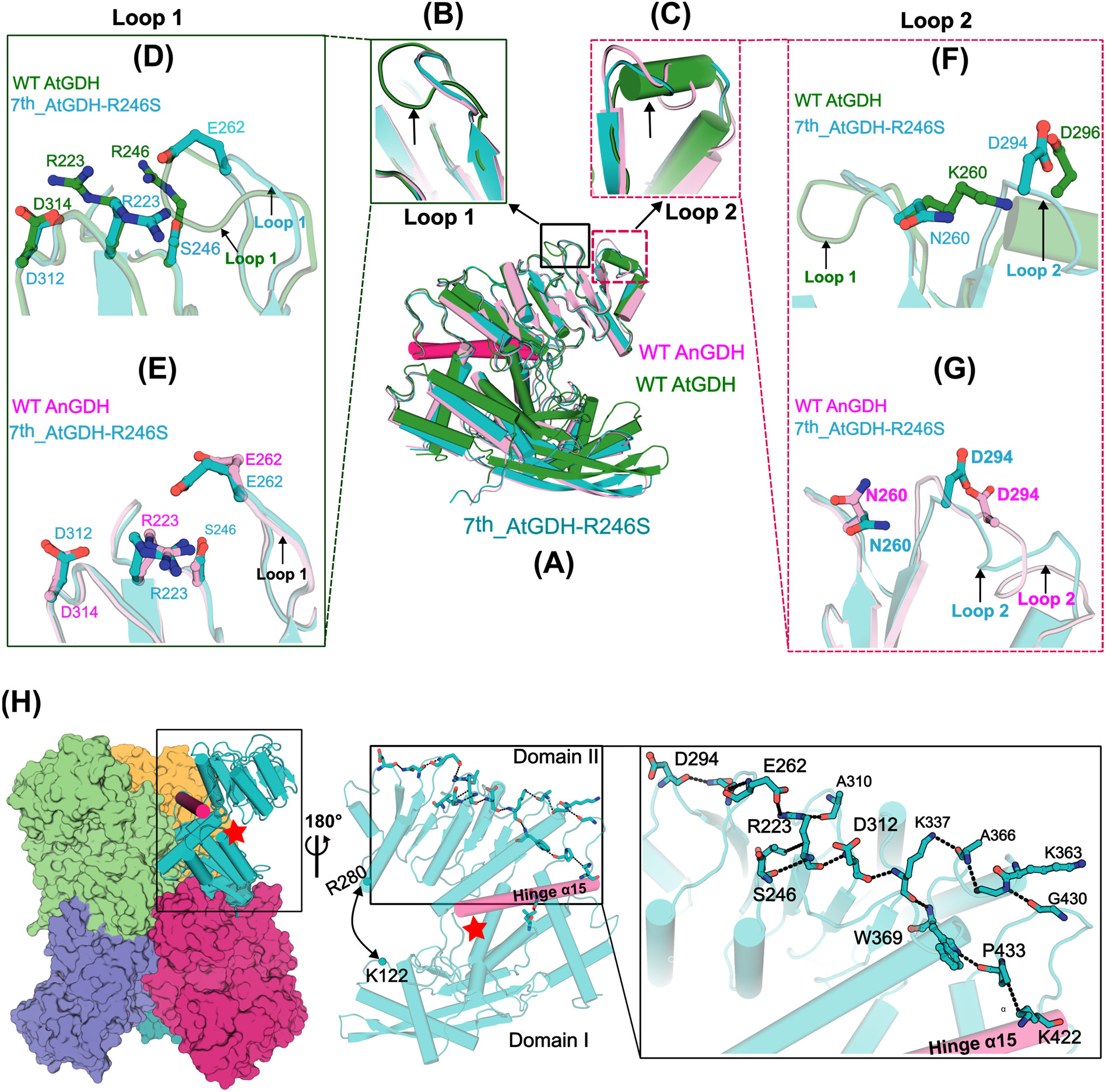
Altered network in Domain II of 7^th^_AtGDH-R246S mutant. (**A**) Superposed structures of 7^th^_AtGDH- R246S (teal carbon) with the wild type AtGDH (green carbon) and AnGDH (pink carbon) are shown. Insets highlight the structural fold differences in Loop 1 **(B)** and Loop 2 **(C)** of these structures. A zoom-in view of Loop 1 shows the differences in the orientation of conserved R223 and the residues of Loop 1 in the 7^th^_AtGDH-R246S mutant structure in comparison to WT AtGDH **(D)** and WT AnGDH **(E)**. **(F)** Superposed structures of 7^th^_AtGDH-R246S and WT AtGDH show the differences in the orientation of N260 (K260 in WT AtGDH) and the displaced D294 (D296 in WT AtGDH). **(G)** The flipped N260 residue in 7^th^_AtGDH-R246S and WT AnGDH are shown. Interaction of N260 with T297 (K299 in WT AnGDH) and D294 (N296 in WT AnGDH) are shown. In WT AnGDH, the N260 mediated interactions are missing. **(H)** Surface representation of 7^th^_AtGDH-R246S mutant hexamer. The inset shows a continuous interacting network in Domain II for one subunit of the 7^th^_AtGDH-R246S mutant structure. The interactions (shown as dashed lines) between the residues (ball and sticks) range from the tip of the helix α11 to the connecting loop close to the hinge helix α15 (shown as a pink ribbon) in the 7^th^_AtGDH-R246S mutant. 180° rotated zoom-in view of the interacting network of the 7^th^_AtGDH-R246S mutant is shown. The star symbol represents the active site.

Overall, the substitutions near Loop 1 region, particularly R246 to S perturbes the whole network around the tip region of Domain II. Consequently, it is manifested in a wider mouth opening observed in the 7^th^_AtGDH-R246S mutant structure. Together, the 7^th^_AtGDH-R246S mutant structure confirms the correlation between the extended mouth openings in open conformation with the gain in substrate-dependent cooperativity.

## 4.4 Discussion

GDH is an essential enzyme that interlinks catabolic and biosynthetic pathways in all living systems. Two homologous enzymes-AtGDH and AnGDH, despite sharing high sequence identity and structural fold, show disparities in the kinetics of substrate utilization during the catalytic reaction. We have performed extensive structural and biochemical studies to delineate the mysteries related to intrinsic protein dynamics and the impact of remotely located substitutions on conformational dynamics and kinetic properties of GDHs.

### Intrinsic dynamic nature of AtGDH and AnGDH

AtGDH and AnGDH exhibit different kinetic behavior towards their substrate α-ketoglutarate, although both the enzymes have almost identical polypeptide sequences. AtGDH and AnGDH were purified as recombinant enzymes, and the structures were determined using X-ray crystallography and cryo-EM to understand the molecular switch that regulates the kinetic properties of these enzymes. AnGDH and AtGDH are hexameric enzymes with a canonical tertiary structural fold like other hexameric GDHs (Baker et al., 1992; Werner et al., 2005; Dimovasili et al., 2021). Only one previous report presented the structural evidences using X-ray crystallography and MD simulation indicating the presence of heterogeneous opening and closing of the active site in the apo form of a *Thermococcus profundus* GDH (TpGDH) (PDB ID: 1EUZ) (Oroguchi & Nakasako, 2016). However, the heterogeneous states of the subunits of TpGDH showed only minute conformational differences in the opening of the active site cleft (≤1 Å), with no change in the interfacial interactions. Moreover, no significantly distinct quaternary structural ensembles of hexameric GDH has yet been reported. So far, none of the previous studies on the GDHs has correlated the structural dynamics with the kinetic properties of this enzyme.

Our structural data on AtGDH (AtGDH-I, II, and III ensembles) are the first direct evidence of an apo-GDH captured in the different conformational sub-states. Additionally, we report the first closed conformations (mouth opening, ∼8 Å) of the apo-forms of AtGDH (AtGDH-III-em) and AnGDH (AnGDH-em). Hence, these structural data from AnGDH and AtGDH indicate that GDHs are intrinsically dynamic proteins and that structural dynamics play an important role in kinetic regulation.

### Amino acid substitutions, located remotely from the active site, alter the Domain II dynamics and kinetic properties in AtGDH and AnGDH

In the enzymes, the structure and its dynamics are linked to their catalytic properties (Torgeson et al., 2022). Structural analysis comparing AnGDH and AtGDH ensembles reveals similarities in their closed quaternary structures, while distinctions in quaternary structures bearing two conformational states (AnGDH:5XVI and AtGDH-I structures). PSN analysis indicated that the major substitutions in Domain II modifies the interacting networks extending from the hinge helix to the Domain II tip in these homologous enzymes. Consequently, these altered networks are manifested in the degree of mouth opening with an observable difference of ∼4 Å between AtGDH (∼17 Å) and AnGDH (∼21 Å). Simultaneously, it influences the inter-subunit contacts between them. Previously, the wide-open catalytic domain leading to more inter-subunit interactions has been rationalized for the positive cooperative behavior for pyruvate binding in homo-tetrameric lactate dehydrogenase from *E. coli* and *F. nucleatum*. The narrow catalytic domain opening in P. aeruginosa with fewer inter-subunit contacts but relatively favorable conformation for substrate binding exhibited Michaelis kinetics (Furukawa et al., 2018). Other studies on homologous monomeric enzymes - glucokinase (allosteric enzyme) and hexokinase I (following Michaelis-Menten kinetics) have demonstrated the variation in mouth opening and conformational dynamics to contribute towards differential kinetic properties for the same substrate saturation (Kamata et al., 2004; Larion & Miller, 2012). Therefore, the variation in domain dynamics and the extent of mouth opening might be considered as one of the primary reasons for the differential kinetic behavior towards AKG substrate in AnGDH and AtGDH. To obtain direct molecular evidence, we attempted to introduce AKG-dependent cooperativity in AtGDH, a Michaelias enzyme, based on the rationale from structural analysis.

Multiple mutagenesis were performed across various regions of the non-allosteric AtGDH, yet only the R246S mutation induced cooperative behavior. Conversely, this mutation (S246 to R) caused a decrease in cooperativity in AnGDH. The R246 is located approximately ∼15–20 Å away from the active site, yet the kinetic parameters of the AtGDH mutant were affected. Thus, our studies indicate that the residues located remotely on Domain II and the active site are wired and functionally synchronized. Previously, the influence of the remote residues (residing ∼15-20 Å away from the active site) on the catalytic activity has been reported in dihydrofolate reductase (DHFR) (Brown et al., 1993; Dion et al., 1993; Cameron & Benkovic, 1997) and LovD proteins (Jimenez-Oses et al., 2014). Numerous studies demonstrate that surface and active site residues are connected by a network that serves as an energy transfer pathway to facilitate enzyme catalysis (Agarwal et al., 2019). Further, the role of remote substitution at R246S was rationalized by the 7^th^_AtGDH-R246S mutant structure that acquired a wider mouth opening compared to the WT AtGDH. In WT AtGDH, the R246 displaces another conserved R223 in the vicinity, disrupting the interaction with Loop 1. However, it is substituted to S246 in AnGDH and thus favors the conserved R223 to orient towards Loop 1 and form a continuous network from hinge to Domain II tip (Fig. 3F). Similarly, the 7^th^_AtGDH-R246S mutant has conserved R223, acquiring the same orientation as observed for the WT AnGDH. The R246S substitution interacts with E262 from the shortened Loop 1, thereby pulling the Loop 2 closer and leading to a comparatively wider mouth opening. However, a similar observation is absent in the 2^nd^_AtGDH-ΔT262-A263 mutant structure due to the presence of both R246 and R223 near the mutated Loop 1. Therefore, our studies reveal the crucial role of S246 in conformational dynamics and allosteric behavior in AnGDH. Hence, this study suggests that the varied mouth opening is directly related to discrepancies in kinetic properties of the allosteric AtGDH and non-allosteric AtGDH.

Taken together with our structural and enzyme kinetics data, it is apparent that the wider mouth opening in AnGDH enables a greater number of inter-subunit interactions at the trimeric center as compared to AtGDH (Fig. 8). Consequently, the open conformation becomes the most stable state of AnGDH. In the presence of substrate, the equilibrium shift from the open to partially closed conformation in AnGDH involves intense structural rearrangement, making it a slower enzyme at lower substrate concentration. In contrast, the multiple conformations of AtGDH coexist in equilibrium, and slight structural modifications are required for transitioning between these conformational sub-states. Thus rendering it a highly dynamic and fast enzyme. In general, the findings of our studies emphasize the role of remote amino acid substitutions that can alter the inherent protein dynamics and conformational dynamics leading to disparities in kinetic behavior among closely related homologous enzymes.

**Figure 8.**
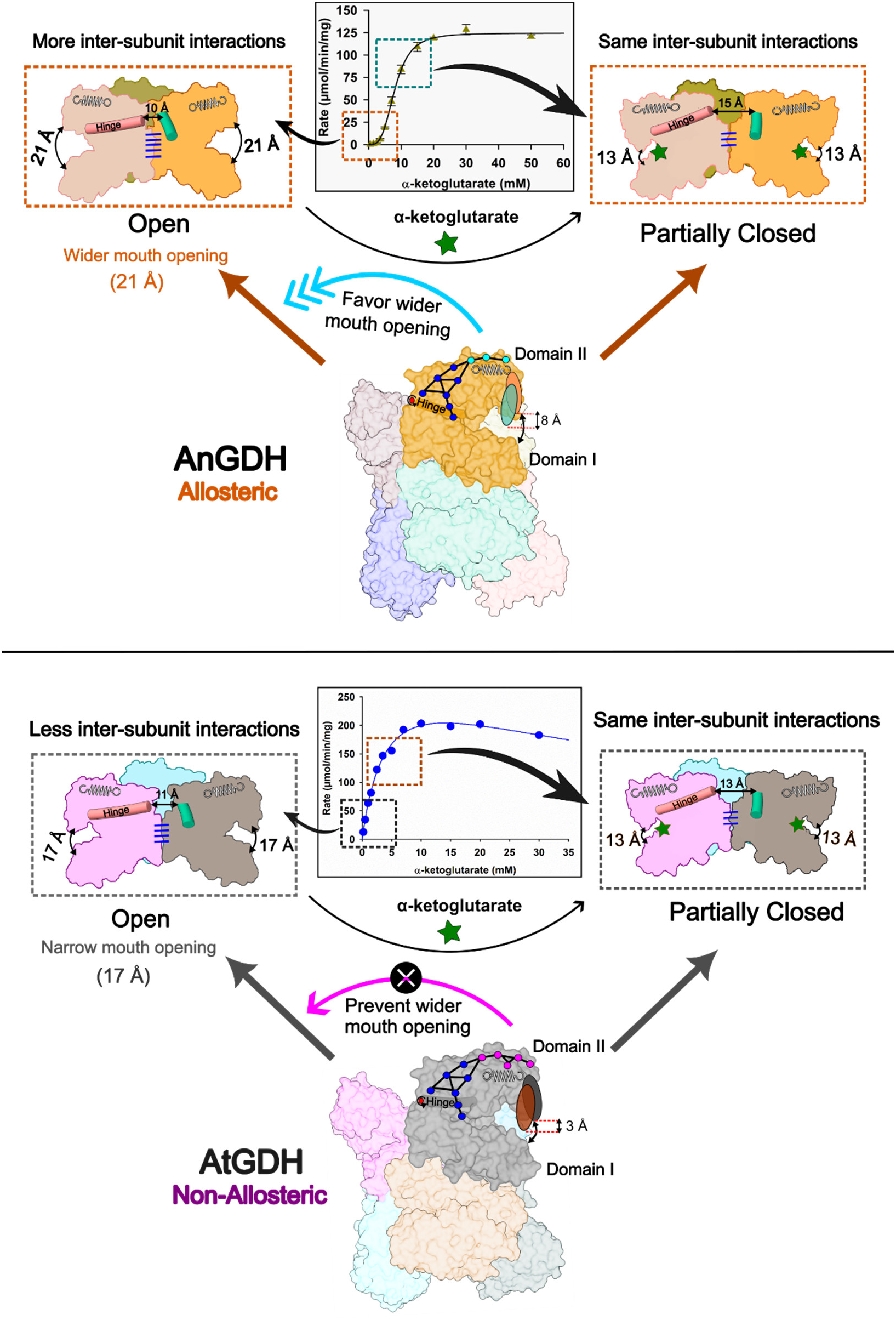
Schematic diagram depicting the relationship between the active site mouth opening and regulation of kinetic properties. The interacting network extending from the active site and hinge helix to Domain II tip is shown as round bead with connecting lines. The blue beads represent conserved residues while the substituted residues in the network which act as spring are shown as cyan beads in AnGDH while pink beads in AtGDH. In allosteric AnGDH, the substituted residues in network (cyan beads) favour wider mouth opening (∼21 Å) and in turn more hydrophobic and electrostatic interactions (shown as blue dash between subunits) at the trimeric inter-subunit region. On the contrary, the substituted residues in network (pink beads) of AtGDH favors narrow opening (∼17 Å), leading to reduction in the number of inter-subunit contacts (blue dash between subunits). The calculated distances (Å) between the adjacent hinge helices are shown by two-headed arrow. The substrate α-ketoglutarate (AKG, green star) bound partially closed conformations of AnGDH and AtGDH (right side panels to graph) have equivalent no. of inter-subunit contacts. On transition from open to partially closed conformations, AnGDH undergoes more loss of interactions than AtGDH. The α-ketoglutarate saturation kinetic plot depicts the population distribution of different enzyme conformations (dashed boxes) at varying substrate concentration. At low substrate concentrations and lower enzyme activity (orange dashed box in graph) are populated by AnGDH enzyme with wider mouth opening (left side of the graph). In contrast, AtGDH enzyme with catalytically competent conformation for AKG binding (left side of the graph) shows higher activity even at lower substrate concentrations (black dashed box in graph). At higher substrate concentrations, the partially closed conformations of AnGDH and AtGDH can be considered of having equivalent activity profile (green dashed box in AnGDH and blue dashed box in AtGDH, and conformers shown on the right side of respective graphs).

## Acknowledgements

We thank Dr. Ravindra Makde and Dr. Biplab Ghosh at the PX-BL21 beamline (BARC) at Indus-2, RRCAT, Indore, India for their support in diffraction data collection. We acknowledge the “Protein Crystallography Facility” at IIT Bombay. We also thank Dr. Vinothkumar Kutti Ragunath and Dr. Sucharita Bose for cryo-EM data collection at the “National Electron Cryo-Microscopy Facility”, Institute for Stem Cell Science and Regenerative Medicine (inStem), Bengaluru, India. Generous computational support from the “SERB and IoE-sponsored National Facility for cryogenic Transmission Electron Microscopy” at IIT Bombay for cryo-EM data processing is also acknowledged. We would like to thank “Spacetime High-Performance Computing (HPC)” resource at IIT Bombay, Mumbai, India for generous computing time. Fellowships to BKJG from Department of Biotechnology (DBT), Ministry of Science and Technology, India and to PD from Council of Scientific & Industrial Research (CSIR) are also acknowledged. The work was supported by research funding to PB from Science and Engineering Research Board (SERB), Govt. of India (Grant No. CRG/2021/002404).

## Author contributions

PB conceived the idea, obtained funding and coordinated the study. BKJG purified the proteins, performed enzyme assays and determined the crystal structures under the supervision of PB. The enzyme kinetics experiments were performed by BKJG with suggestions from NSP under the supervision of PB. PD set up the computational facility and softwares for cryo-EM data processing in PB group. BKJG and PD processed the cryo-EM datasets with the inputs from SK under the supervision of PB. BKJG refined and analyzed the cryo-EM structures with inputs from PD and SK under the supervision of PB. AS performed MD simulation and analyzed the trajectories. PKM purified and performed enzyme kinetics of two AnGDH mutants with inputs from BKJG. BKJG and PB wrote the manuscript with inputs from all authors.

## Supplementary Materials

### Supplementary movie legends

**Supplementary movie 1**

Conformational heterogeneity in the cryo-EM electron density map of non-allosteric apo-AtGDH. Two predominant ensembles of partially closed and closed conformations are observed. The dynamic nature of Domain II in AtGDH is captured in the cryo-EM structures.

**Supplementary movie 2**

Domain II dynamics of non-allosteric AtGDH. The movie displays the extent of mouth opening in AtGDH and the role of residues (shown as sticks) in regulating the motion of Domain II.

**Supplementary movie 3**

Movement of Domain II in allosteric AnGDH. Movie shows the wider mouth opening in AnGDH as compared to AtGDH.The residues in Domain II influencing the extent of mouth opening are shown as sticks.

**Table S1.**
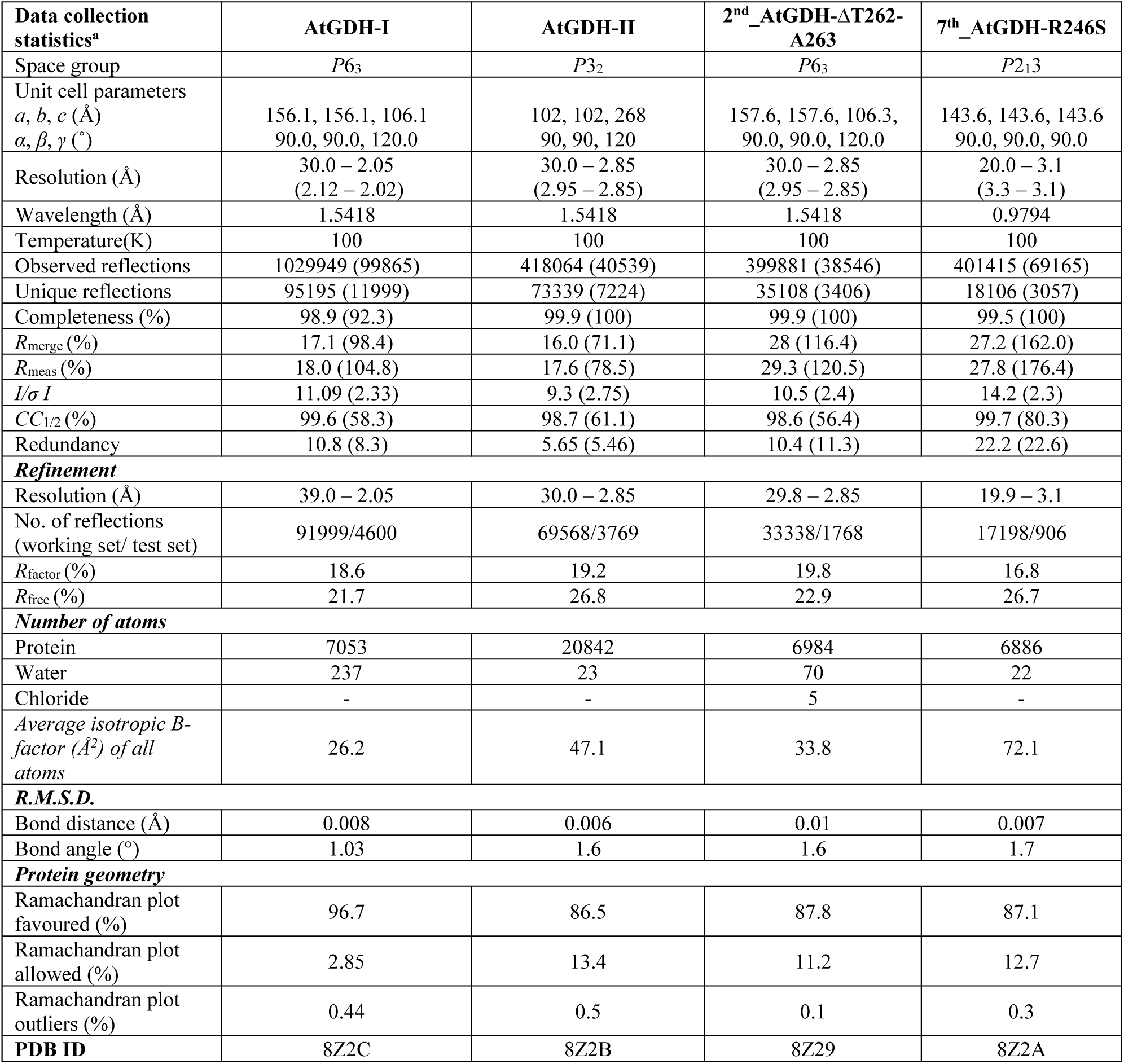
Data collection and refinement statistics of AtGDH crystal structures.

**Figure S1.**
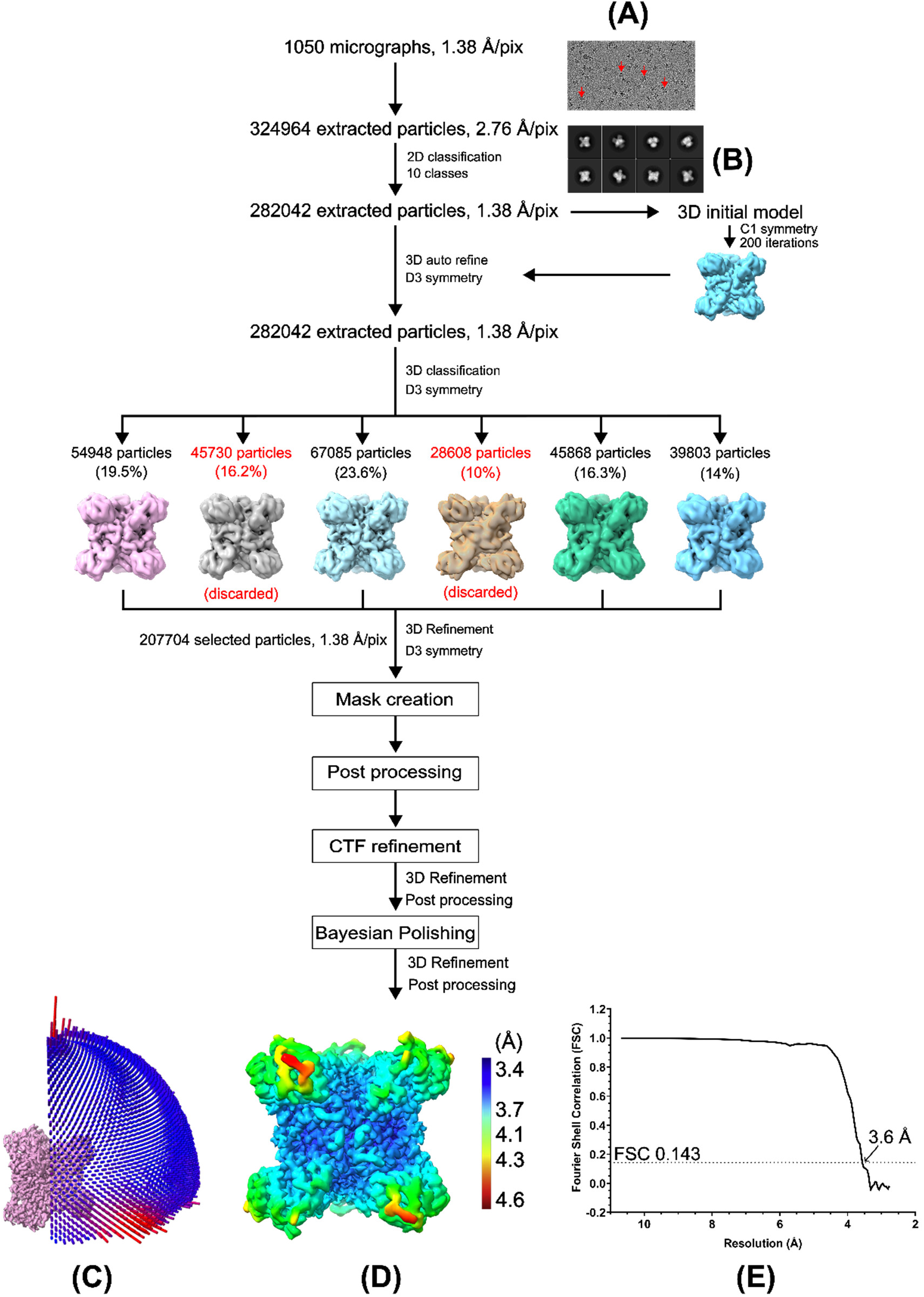
Workflow for cryo-EM data processing of AnGDH. A stepwise method display the particle extraction, classification (2D and 3D) and refinement considered for data processing. (**A**) A representative micrograph collected for AnGDH specimen. The red arrow indicates the AnGDH particles (‘X’ or clove leaf shaped black spots) present on the micrographs. (**B**) Representative 2D class averages of AnGDH particle images are shown. **(C)** Angular distribution of particles used for reconstruction of AnGDH map. **(D)** Final reconstructed map of AnGDH colored by local resolution determined using Relion v3.0. The color scale display the resolution range. **(E)** The Fourier Shell correlation (FSC) of reconstruction at gold standard (0.143) and the value is marked in the plot.

**Figure S2.**
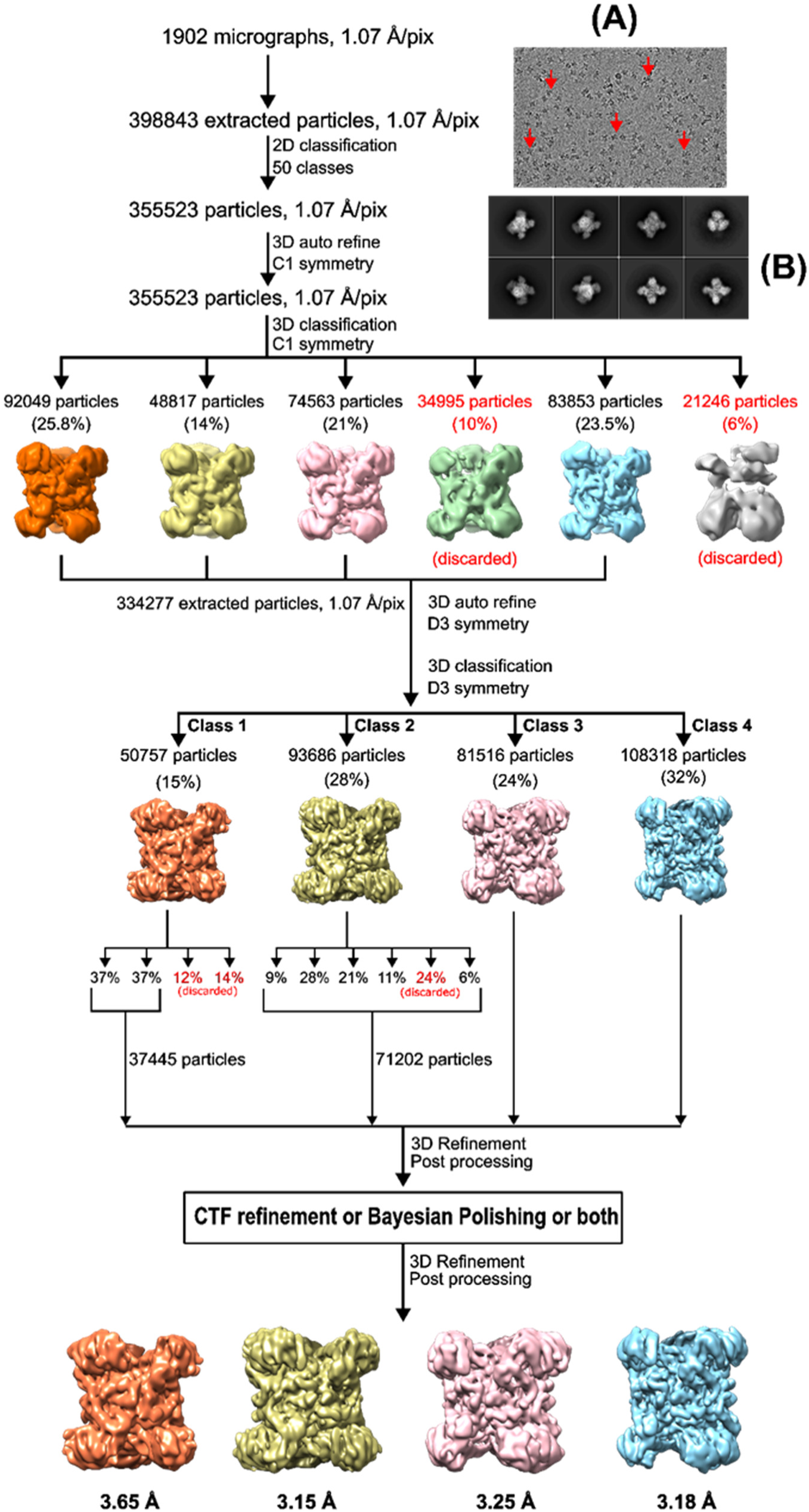
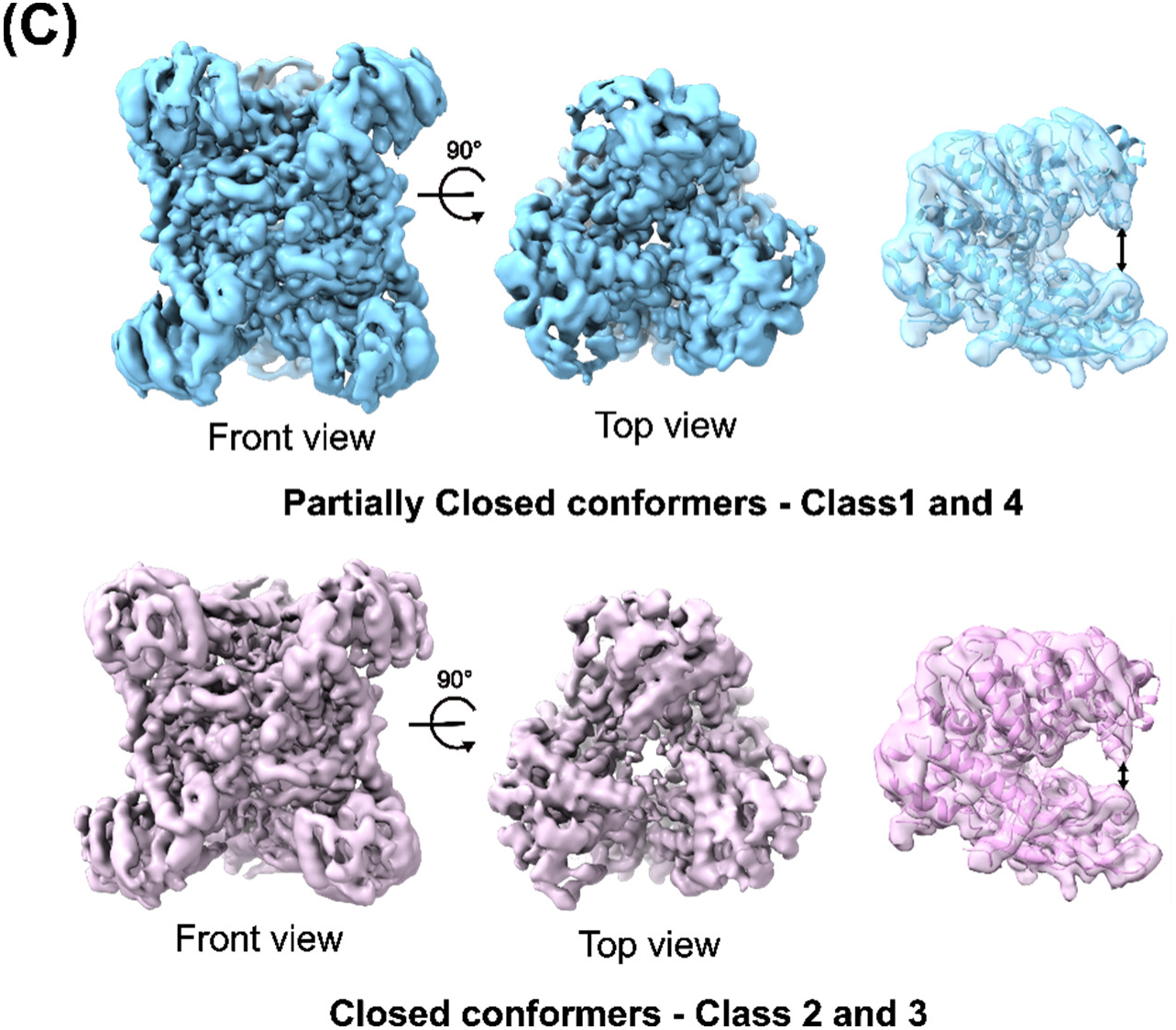
Cryo-EM data processing workflow of AtGDH. The stepwise method consisting of particle picking, classification (2D and 3D), and refinement strategies used for obtaining the reconstructed map is shown. **(A)** A representative cryo-EM micrograph of AtGDH specimen. The black ‘X’ or clove leaf shaped spots (marked by red arrow) are the AtGDH particles images on the collected micrographs**. (B)** Representative 2D class averaging of AtGDH particles images in multiple orientations are shown. The four final reconstructed maps of AtGDH particles with the predicted resolution (FSC 0.143) gold standard are reported. **(C)** Front and top views of the maps are presented for the different classes. Right panels (for both conformers) with a model fitted to one subunit of the map. The double-sided arrow depicts the mouth opening.

**Figure S3.**
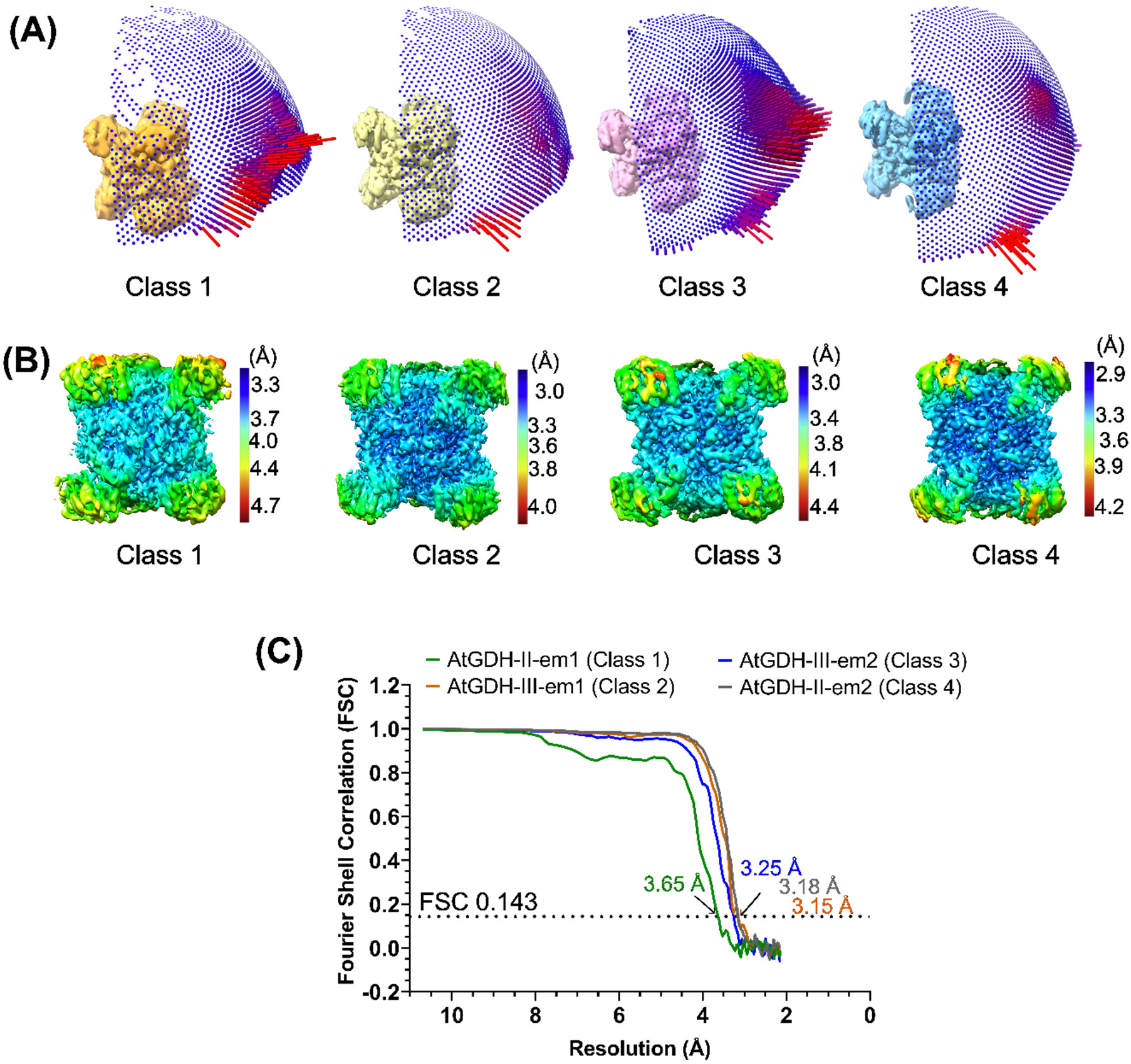
Analysis of cryo-EM map of AtGDH ensembles. (**A**) The angular distribution of particles used for reconstruction of each classes 1, 2, 3 and 4 are shown. **(B)** Local resolution maps calculated using Relion v3.0 are shown for all four classes. A five-color code is used to color the reconstructed map on the basis of resolution at different parts. **(C)** A gold standard FSC 0.143 curves displays the resolution of four reconstructed maps of AtGDH (AtGDH-II-em1, AtGDH-II-em2, AtGDH-III-em1, and AtGDH-III-em2).

**Table S2.**
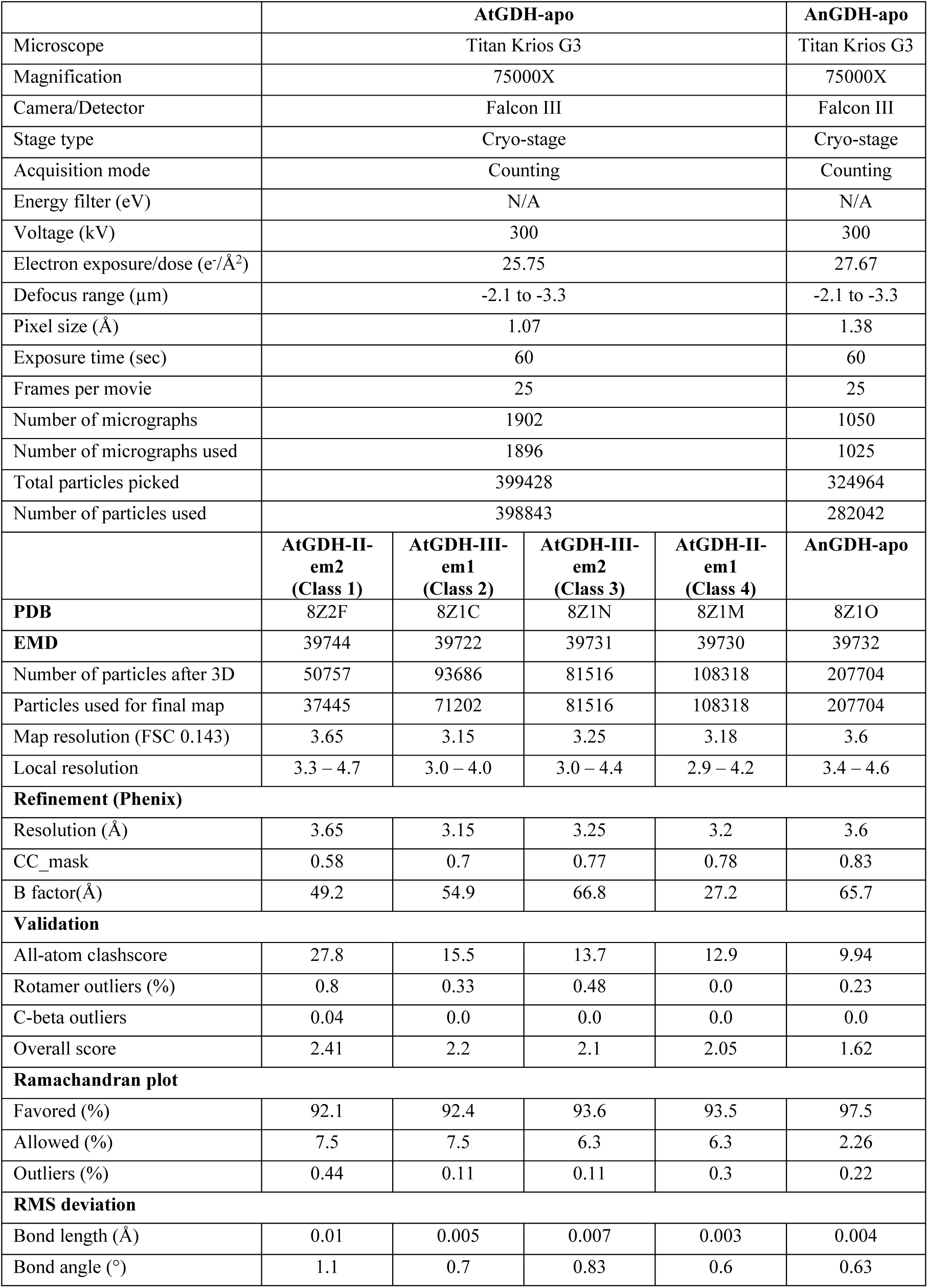
cryo-EM data collection and refinement statistics.

|  | **AtGDH-apo** |  |  |  | **AnGDH-apo** |
| --- | --- | --- | --- | --- | --- |
| Microscope | Titan Krios G3 |  |  |  | Titan Krios G3 |
| Magnification | 75000X |  |  |  | 75000X |
| Camera/Detector | Falcon III |  |  |  | Falcon III |
| Stage type | Cryo-stage |  |  |  | Cryo-stage |
| Acquisition mode | Counting |  |  |  | Counting |
| Energy filter (eV) | N/A |  |  |  | N/A |
| Voltage (kV) | 300 |  |  |  | 300 |
| Electron exposure/dose (e<sup>-</sup>/Å<sup>2</sup>) | 25.75 |  |  |  | 27.67 |
| Defocus range (μm) | -2.1 to -3.3 |  |  |  | -2.1 to -3.3 |
| Pixel size (Å) | 1.07 |  |  |  | 1.38 |
| Exposure time (sec) | 60 |  |  |  | 60 |
| Frames per movie | 25 |  |  |  | 25 |
| Number of micrographs | 1902 |  |  |  | 1050 |
| Number of micrographs used | 1896 |  |  |  | 1025 |
| Total particles picked | 399428 |  |  |  | 324964 |
| Number of particles used | 398843 |  |  |  | 282042 |
|  | **AtGDH-II-em2**<br>**(Class 1)** | **AtGDH-III-em1**<br>**(Class 2)** | **AtGDH-III-em2**<br>**(Class 3)** | **AtGDH-II-em1**<br>**(Class 4)** | **AnGDH-apo** |
| **PDB** | 8Z2F | 8Z1C | 8Z1N | 8Z1M | 8Z1O |
| **EMD** | 39744 | 39722 | 39731 | 39730 | 39732 |
| Number of particles after 3D | 50757 | 93686 | 81516 | 108318 | 207704 |
| Particles used for final map | 37445 | 71202 | 81516 | 108318 | 207704 |
| Map resolution (FSC 0.143) | 3.65 | 3.15 | 3.25 | 3.18 | 3.6 |
| Local resolution | 3.3 – 4.7 | 3.0 – 4.0 | 3.0 – 4.4 | 2.9 – 4.2 | 3.4 – 4.6 |
| **Refinement (Phenix)** |  |  |  |  |  |
| Resolution (Å) | 3.65 | 3.15 | 3.25 | 3.2 | 3.6 |
| CC_mask | 0.58 | 0.7 | 0.77 | 0.78 | 0.83 |
| B factor(Å) | 49.2 | 54.9 | 66.8 | 27.2 | 65.7 |
| **Validation** |  |  |  |  |  |
| All-atom clashscore | 27.8 | 15.5 | 13.7 | 12.9 | 9.94 |
| Rotamer outliers (%) | 0.8 | 0.33 | 0.48 | 0.0 | 0.23 |
| C-beta outliers | 0.04 | 0.0 | 0.0 | 0.0 | 0.0 |
| Overall score | 2.41 | 2.2 | 2.1 | 2.05 | 1.62 |
| **Ramachandran plot** |  |  |  |  |  |
| Favored (%) | 92.1 | 92.4 | 93.6 | 93.5 | 97.5 |
| Allowed (%) | 7.5 | 7.5 | 6.3 | 6.3 | 2.26 |
| Outliers (%) | 0.44 | 0.11 | 0.11 | 0.3 | 0.22 |
| **RMS deviation** |  |  |  |  |  |
| Bond length (Å) | 0.01 | 0.005 | 0.007 | 0.003 | 0.004 |
| Bond angle (°) | 1.1 | 0.7 | 0.83 | 0.6 | 0.63 |

**Table S3.** Measured active site cleft opening of AtGDH quaternary structures.

| Subunit | Crystal structures |  | Cryo-EM structures |  |  |  |
| --- | --- | --- | --- | --- | --- | --- |
|  | AtGDH-I<br>(Å) | AtGDH-II<br>(Å) | #AtGDH-II-em1<br>(Å) | #AtGDH-II-em2<br>(Å) | AtGDH-III-em1<br>(Å) | AtGDH-III-em2<br>(Å) |
| A | 13.3 | 12.0 | 13.4 | 12.7 | 7.3 | 8.1 |
| B | 13.3 | 11.6 | 13.8 | 12.6 | 7.7 | 8.5 |
| C | 13.3 | 11.1 | 15.1 | 12.7 | 7.5 | 8.2 |
| D | 17.6 | 11.6 | 13.2 | 12.9 | 7.2 | 8.0 |
| E | 17.6 | 11.6 | 10.2 | 13.2 | 7.7 | 8.4 |
| F | 17.6 | 10.8 | 12.4 | 12.9 | 7.4 | 8.1 |

**Figure S4.**
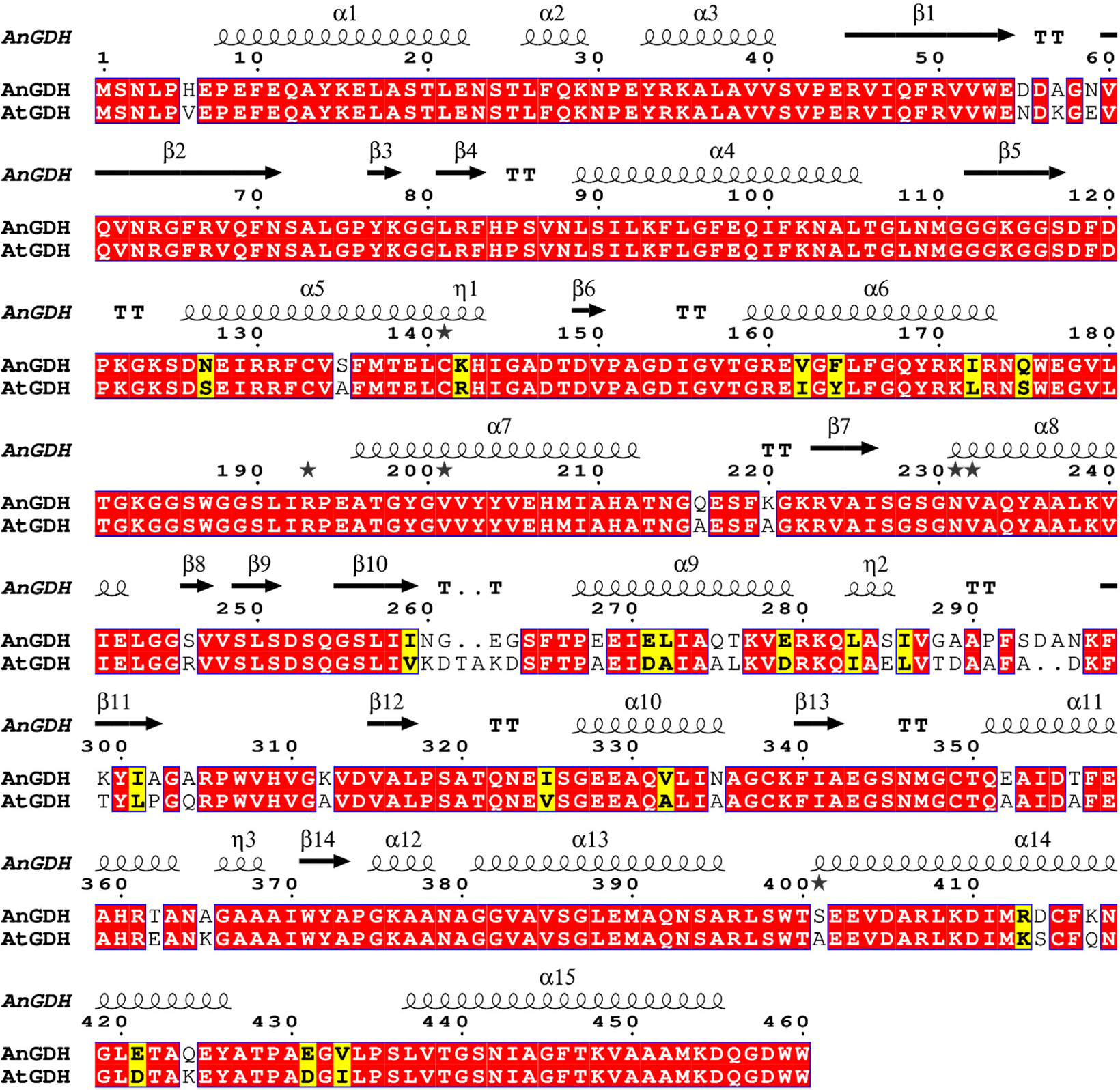
Pairwise-sequence alignment of AnGDH and AtGDH. The red box indicates the conserved amino acid residues. The yellow box indicates the amino acid substitutions with a similar amino acid. The different residues are left unboxed. The missing residues between AnGDH and AtGDH are shown as dots. The predicted secondary structural fold are marked for helices (coil shape), beta-strand (side arrow), and turn (TT). The sequence alignment is generated in ESPript 3.0 (https://espript.ibcp.fr).

**Figure S5.**
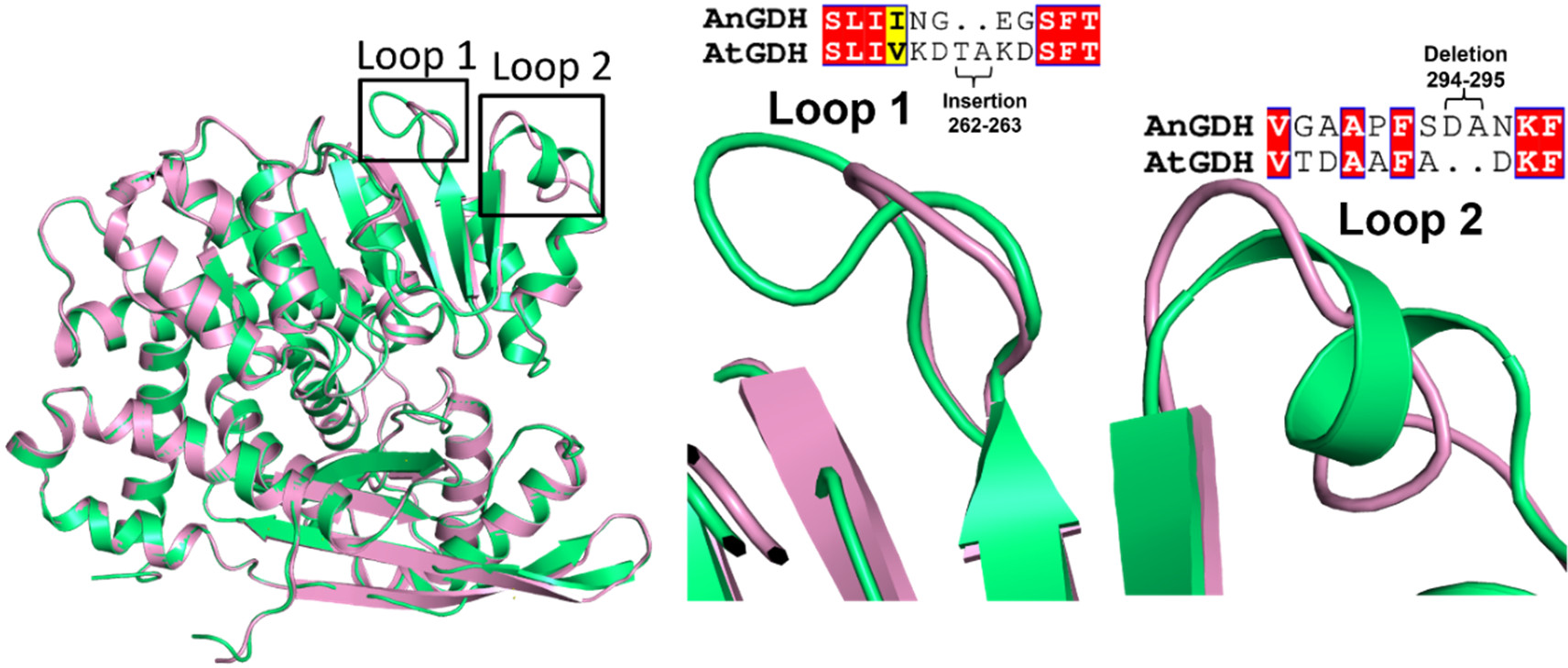
Structural fold differences between AtGDH and AnGDH. Superposed structure of AtGDH (PDB ID: 7ECS, green carbon) and AnGDH (PDB ID: 5XVX, pink carbon). Amino acid substitutions in loop 1 (T262-A263) and deletion in loop 2 (D294-A295) of AtGDH are shown. The sequence alignment on top highlights the amino acid substitutions at these loops.

**Figure. S6.**
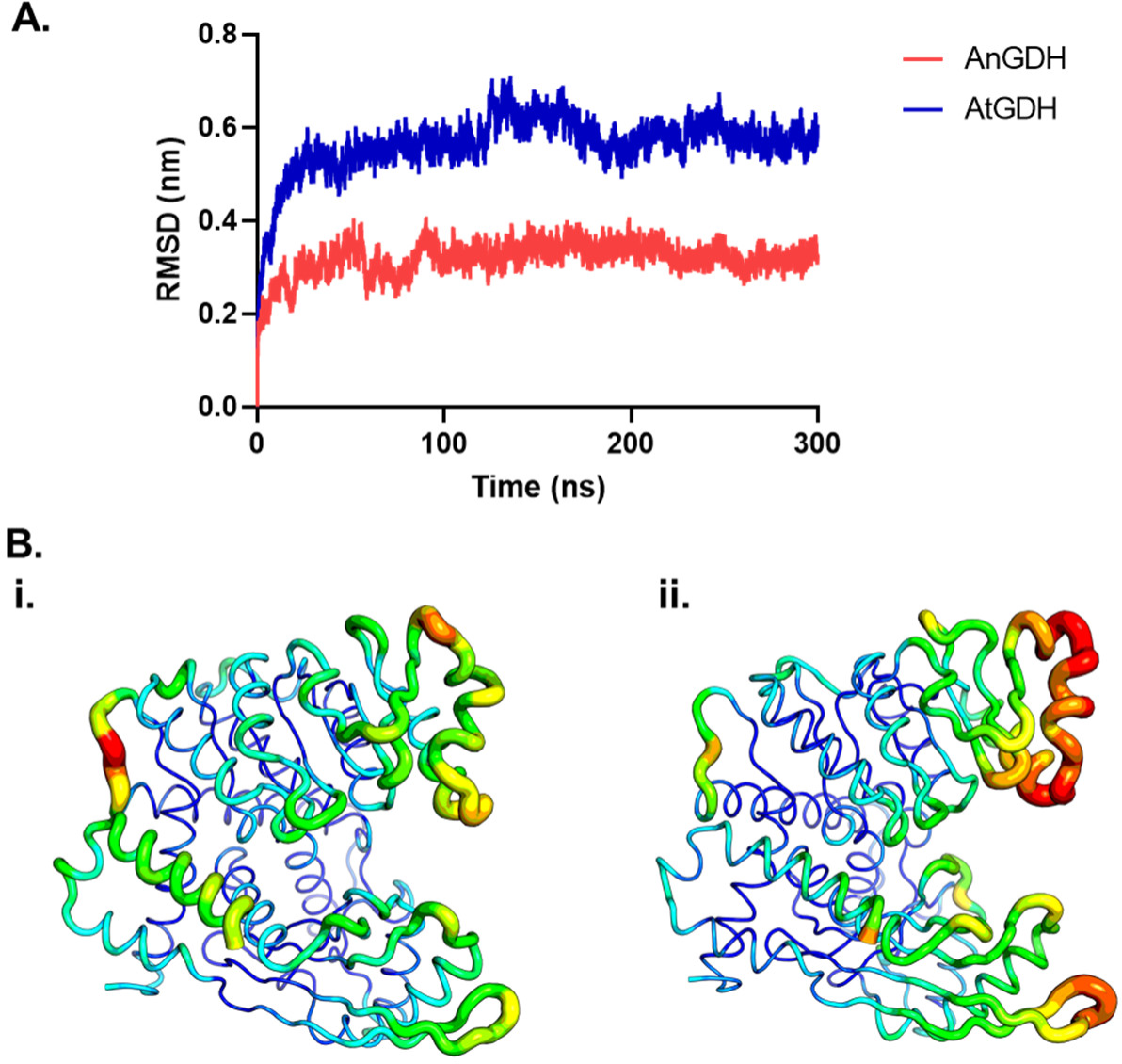
A. Plots of root-mean square deviation (RMSD) of C-alpha atoms of residues of AnGDH (coloured red) and AtGDH (coloured blue) versus simulation time. B. Putty structures of (i) AnGDH and (ii) AtGDH coloured according to the root mean square fluctuations of C-alpha atoms ranging from red (indicating the highest RMSF values) to blue (indicating the lowest RMSF values).

**Figure S7.**
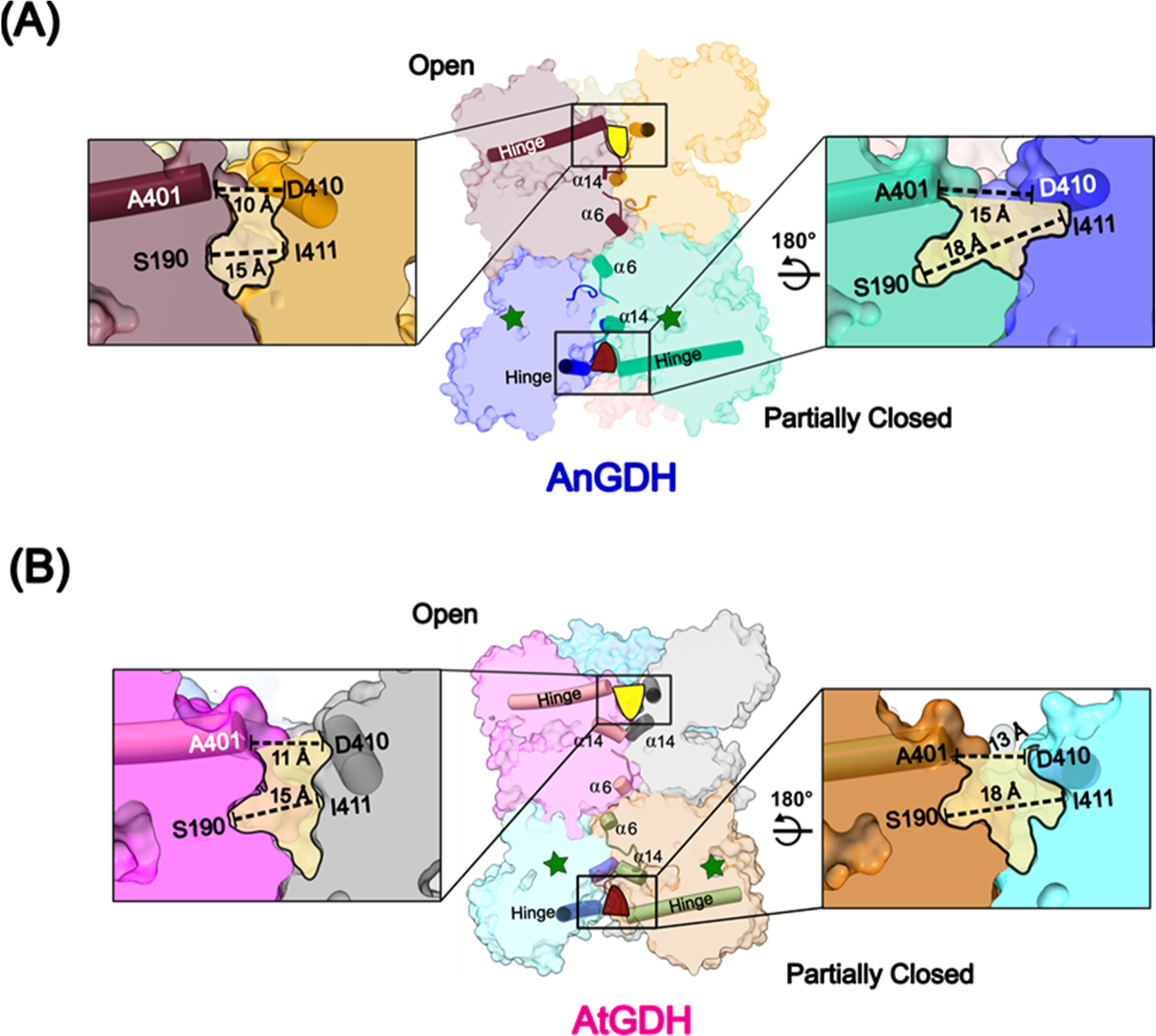
Change in compactness at the trimeric inter-subunit regions of AnGDH and AtGDH. In presence of substrate α-ketoglutarate (green star), transition from open to partially closed conformations occurs. A calculated distances (Å) between the hinge helices of neighboring subunits are shown for the open (left side) and partially closed (right side) subunit pairs of AnGDH (A) and (B). The dashed line highlights the measured distances between the residues S401/A401-D410 and S190-I141 in the open and partially closed conformations of AnGDH and AtGDH.

**Table S4.** Interfacial interactions in AnGDH and different forms of AtGDH quaternary structures. HB: hydrogen bond, SB: salt bridge, PC: partially closed conformation.

| AtGDH | Inter-subunit interactions |  |  |  |  |  |
| --- | --- | --- | --- | --- | --- | --- |
|  | Dimeric |  | Trimeric |  | Diagonal |  |
|  | HB | SB | HB | SB | HB | SB |
| I | 14 | 2 | 16 (PC),<br>17 (open) | 4 (PC and<br>open) | 8 | 4 |
| II | 13 | 3 | 12 | 3 | 6 | 3 |
| II-em1 | 12 | 3 | 9 | 2 | 6 | 2 |
| II-em2 | 20 | 3 | 14 | 5 | 8 | 10 |
| III-em1 | 22 | 3 | 14 | 3 | 9 | 7 |
| III-em2 | 20 | 3 | 11 | 4 | 9 | 9 |
| AnGDH<br>(5XVI) | 28 | 12 | 18 (PC),<br>20 (open) | 6 (PC and<br>9 (open) | 13 | 14 |

**Figure S8.**
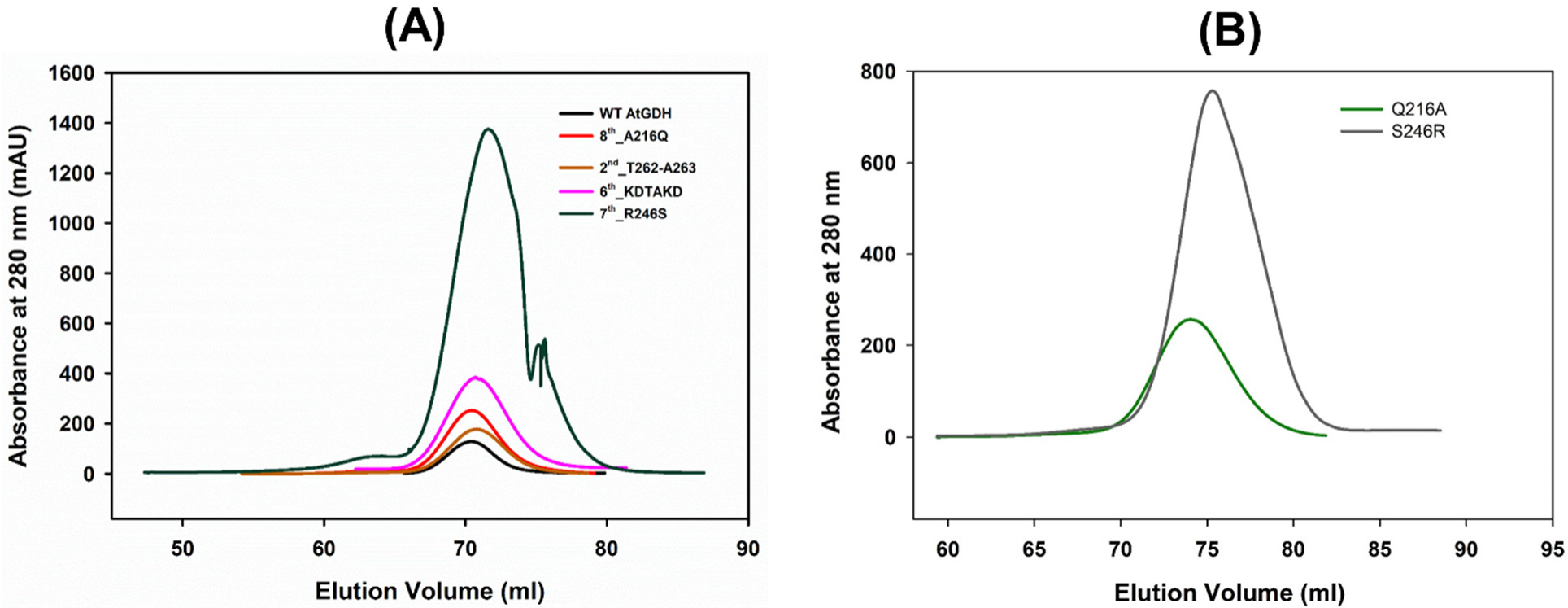
Size exclusion chromatography. Elution profile of AtGDH mutants **(A)** and AnGDH mutants **(B)**.

**Figure S9.**
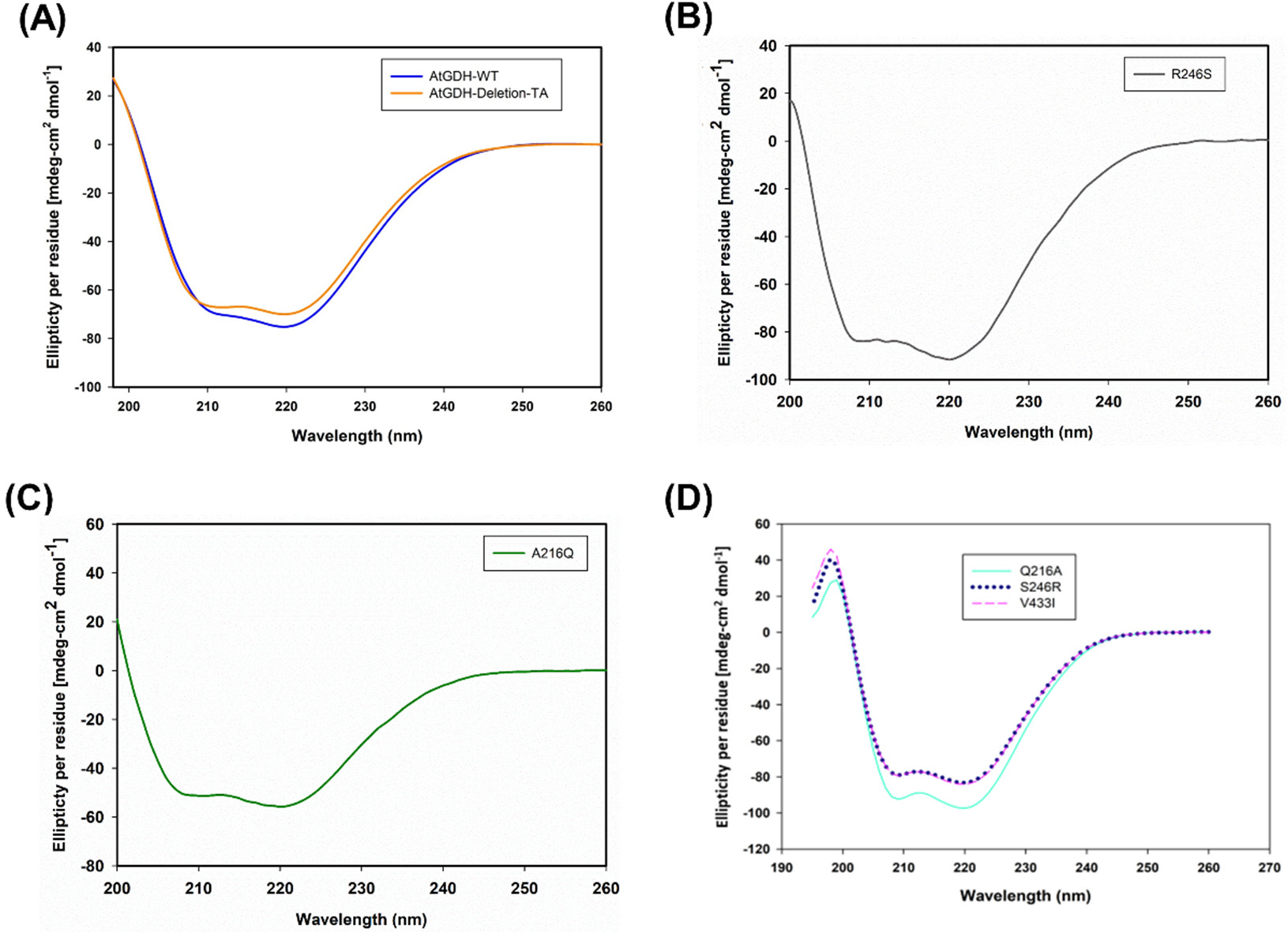
Circular Dichroism (CD) spectrum of AtGDH and AnGDH mutants. (**A**) The spectra absorbance of wild type AtGDH (blue) and 2^nd^_AtGDH-ΔT262-A263 (orange). CD spectra of 7^th^_AtGDH-R246S (**B**) and 8^th^_AtGDH-A216Q (green) (**C**). (**D**) Spectra absorbance of AnGDH mutants – Q216A and S246R in the wavelength range of 198-260 nm are shown.

**Figure S10.**
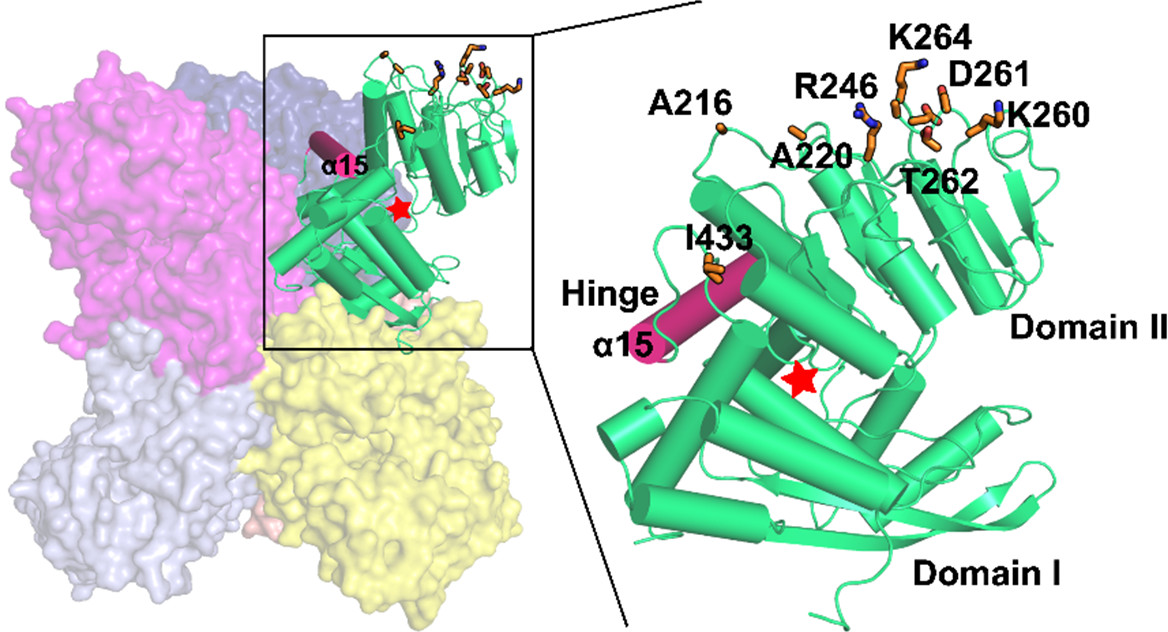
Location of mutated residues in Domain II of AtGDH. The hexameric assembly of AtGDH is shown as surface and cartoon representation. Zoom-in view highlights one subunit with Domain I and Domain II separated by hinge helix (pink cartoon). The location of eight amino acid mutations performed in AtGDH are shown in orange sticks. The active site is shown by a red star symbol.

**Figure S11.**
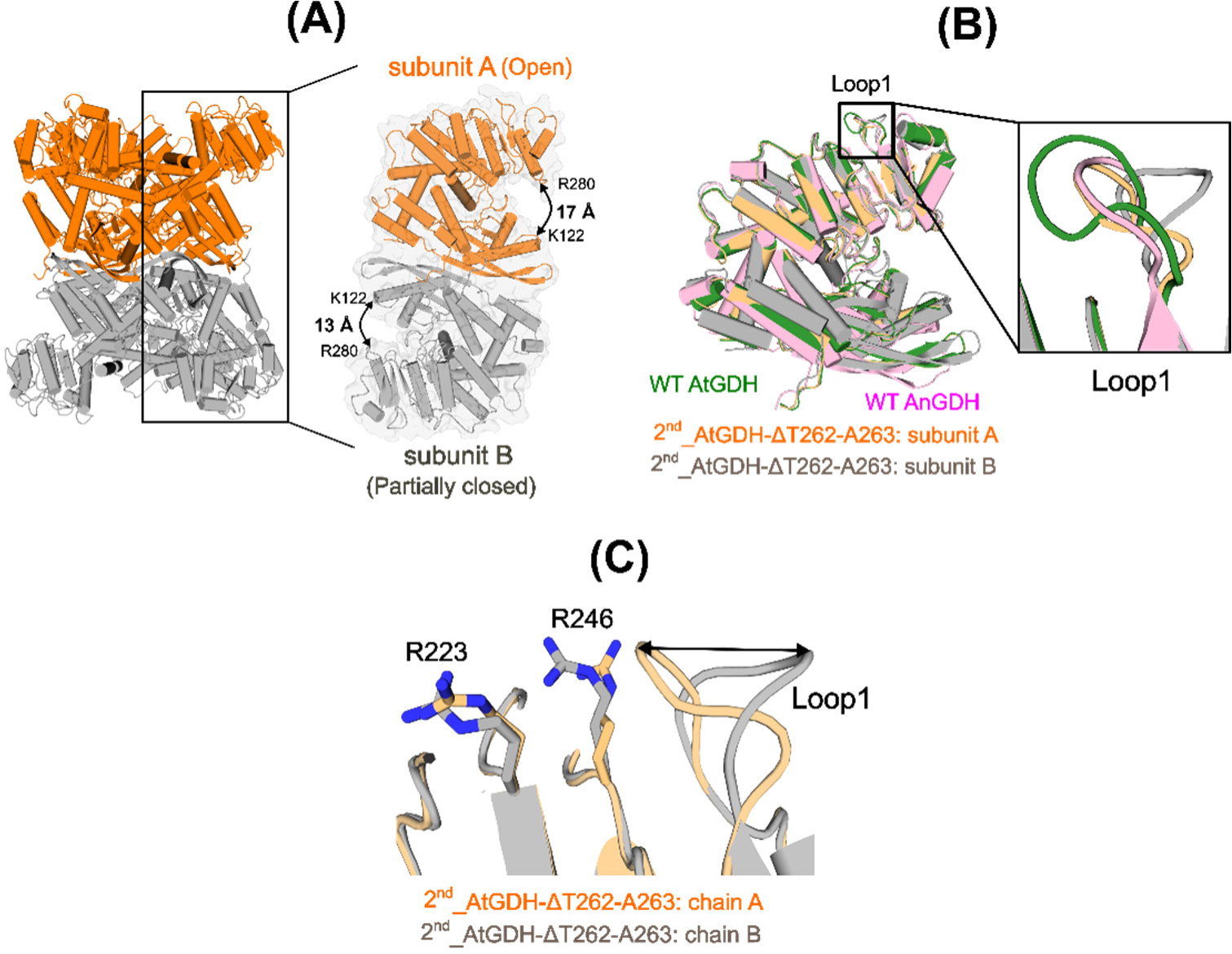
Structural deviations in the 2^nd^_AtGDH-ΔT262-A263 mutant. **(A)** Hexameric structure of 2^nd^_AtGDH-ΔT262-A263 mutant displaying the presence of two co-existing conformations – open (orange) and partially closed (gray). Inset shows the two distinct conformers with the mouth opening distance measured between the active site cleft residues – K122 and R280. **(B**) Superposed structure of 2^nd^_AtGDH-ΔT262-A263 (orange and gray carbon) with the wild type AtGDH (green carbon) and AnGDH (pink carbon). Inset shows the difference in orientation of mutated Loop1 (ΔT262-A263) found in 2^nd^_AtGDH-ΔT262-A263 mutant in comparison to WT AtGDH and WT AnGDH. **(C)** Presence of two distinct conformations of mutated Loop 1 (ΔT262-A263) in the partially closed (gray cartoon) andopen conformations (orange cartoon). The two-sided arrow indicates the Loop1 orientations in the open and partially closed conformers. The structural rearrangement of the neighboring residues - R223 and R246 (sticks with respective) color codes in response to mutated Loop1 (ΔT262-A263) movement are shown.

**Figure S12.**
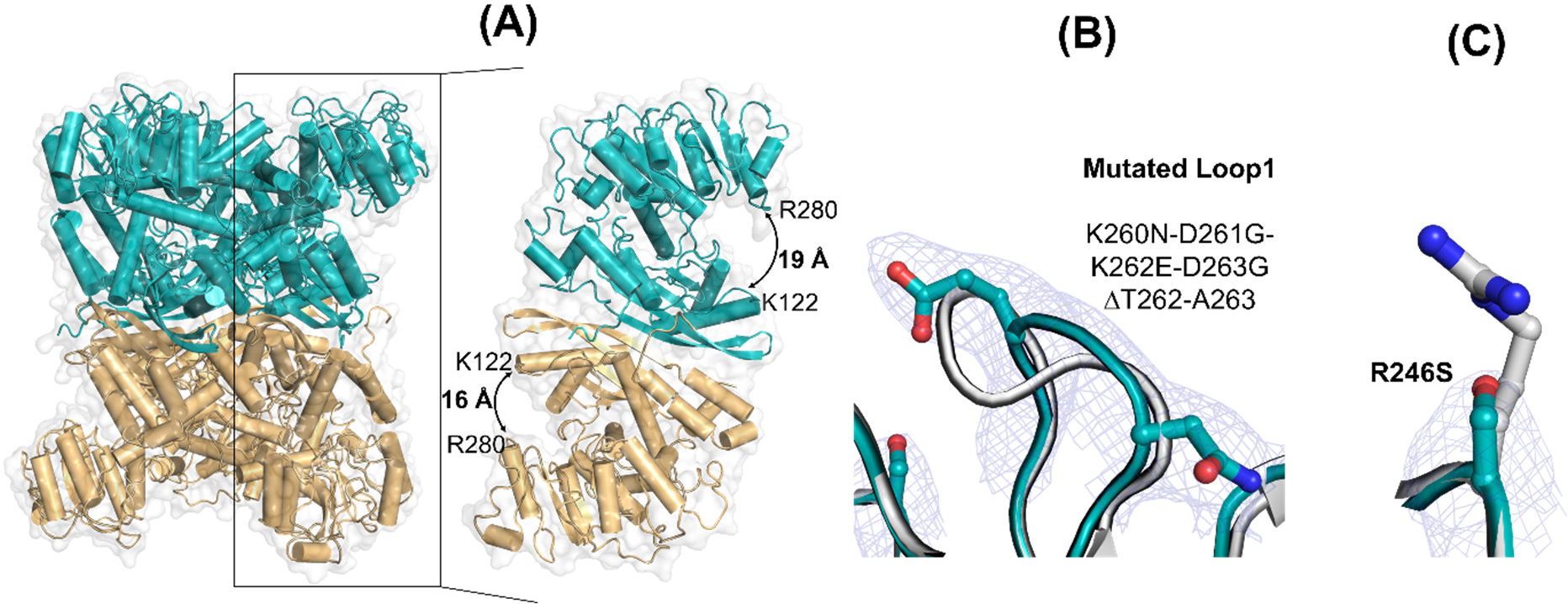
Structural arrangement of the 7^th^_AtGDH-R246S mutant and the supporting electron densities around the mutation point. **(A)** Cartoon representation of the structure of 7^th^_AtGDH-R246S mutant with two conformationally distinct monomers (deep teal and light orange cartoons) stacked on top of each other along the 2-fold axis. The measured active site cleft opening between the cleft residues-K122 (Domain I) and R280 (Domain II) is shown and indicated by an arrow. The electron density map (2*F*_o_-*F*_c_) contoured at 3σ is shown as a blue mesh around the mutated amino acids – Loop 1 (K260N-D261G-K262E-D263G ΔT262-A263 **(B)** and R246S **(C)**. The superposed AtGDH-WT residues are the corresponding positions-Loop 1 and 246 are shown as gray sticks. The mutated residues in the structure of the 7^th^_AtGDH-R246S mutant are shown in deep teal sticks.

**Table S5.**
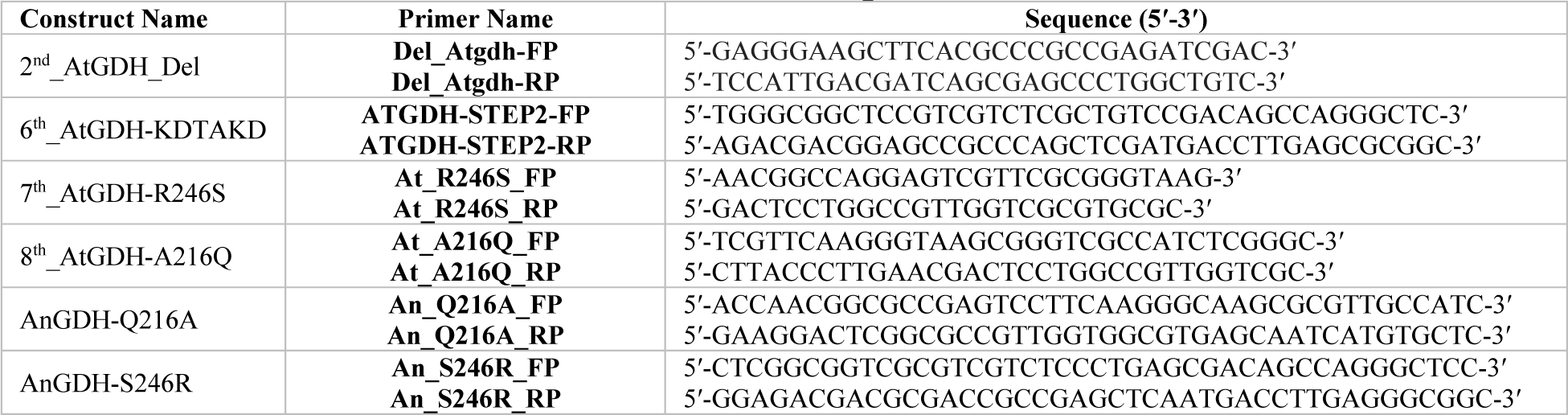
List of primers.

